# Computational Modelling of Resistance to Hormone-Mediated Remission in Childhood Absence Epilepsy

**DOI:** 10.1101/2025.09.04.674333

**Authors:** Maliha Ahmed, Sue Ann Campbell

**Affiliations:** Department of Applied Mathematics and Centre for Theoretical Neuroscience, University of Waterloo, Waterloo, ON, Canada

**Keywords:** childhood absence epilepsy, conductance-based model, remission, thalamocortical, frontocortical

## Abstract

Childhood absence epilepsy (CAE) often resolves during adolescence, a period marked by hormonal and neurosteroid changes associated with puberty. However, remission does not occur in all individuals. To investigate this clinical heterogeneity, we developed a simplified thalamocortical model with a layered cortical structure, using deep-layer intrinsically bursting (IB) neurons to represent frontal cortex and regular spiking (RS) neurons modelling the parietal cortex. By simulating two cortical configurations, we explored how variations in neuronal composition and frontocortical connectivity influence seizure dynamics and the effectiveness of allopregnanolone (ALLO) in resolving pathological spike-wave discharges (SWDs) associated with CAE. While both models exhibited similar physiological and pathological oscillations, only the parietal-dominant network (with a higher proportion of regular spiking neurons in layer 5) recovered from SWDs under increased frontocortical connectivity following ALLO administration. These findings suggest that neuronal composition critically modulates ALLO-mediated resolution of SWDs, providing a mechanistic link between structural connectivity and clinical outcomes in CAE, and highlighting the potential for personalized treatment strategies based on underlying network architecture.

## 1 Introduction

Childhood absence epilepsy (CAE) is a common pediatric epilepsy disorder, accounting for approximately 18% of all childhood epilepsies, with onset typically occurring between the ages of 4 and 12 years (Hirsch et al., 2022). Characterized by brief episodes of impaired consciousness, typical absence seizures present with bilateral synchronous 2.5–5 Hz spike-wave discharges (SWDs) on electroencephalography (EEG) (Hirsch et al., 2022). While antiepileptic drugs are effective for most patients, with approximately 70% achieving remission, some cases progress to more severe forms of epilepsy (Kessler and McGinnis, 2019). Interestingly, circumstantial evidence regarding the prognosis of untreated epilepsy suggests spontaneous remission rates between 31–42%, although these figures are not specific to childhood-onset epilepsy syndromes (Kwan and Sander, 2004; Beghi et al., 2015). Several factors, including the use of antiepileptic drugs, genetic predisposition, and intrinsic connectivity differences, are hypothesized to influence remission; however, understanding of these factors remains limited due to their considerable variability among patients.

The thalamocortical circuit—comprising pyramidal neurons in the cortex, thalamic relay neurons, and thalamic reticular neurons—plays a critical role in both normal 7–10 Hz sleep spindle generation and pathological SWD activity (Sitnikova et al., 2014). Evidence indicates that SWDs may originate from slower oscillations within focal excitable regions, particularly the somatosensory cortex, before being sustained by circuits in the frontal cortex and thalamus (Meeren et al., 2002; Masterton et al., 2013). Notably, children with newly diagnosed CAE who do not respond to treatment demonstrate increased frontocortical connectivity prior to treatment initiation, suggesting fundamental differences in network architecture may influence disease trajectory (Tenney et al., 2018).

Neuronal electrophysiology and intrinsic membrane properties vary significantly based on neuron morphology, cortical layer, and cortical region. Pyramidal neurons in the cortex typically exhibit either regular spiking (RS) or intrinsically bursting (IB) behaviour. Intrinsically bursting neurons tend to have thick apical dendrites as compared to those that are regular spiking, as shown in studies of pyramidal neurons in layers 2/3 and 5 of rat visual cortex (Mason and Larkman, 1990). In slices of sensorimotor and frontal cortex from guinea pigs, regular spiking pyramidals were found in all layers below layer 1, while bursting cells were located mostly in layers 4/5 and deeper (McCormick et al., 1985). Additionally, studies in Sprague-Dawley rats indicate a higher percentage (approximately 73%) of layer 5/6 IB neurons in slices of prefrontal cortex, compared to layer 5 cells in slices of somatosensory (parietal) cortex (approximately 50–60%) (Chagnac-Amitai et al., 1990; Yang et al., 1996). These regional differences in neuronal composition may contribute to the pathophysiological variability observed in CAE.

Given that remission in most cases occurs during adolescence, the association of sex steroid hormones with CAE is suspected and additionally supported by the phenomenon of catamenial seizure exacerbation (van Luijtelaar et al., 2001). Progesterone is one of the key steroid hormones that plays a crucial role in sexual development during puberty. While it is primarily regarded as a female hormone, its significance in males is also recognized (Ghoumari et al., 2014; Guennoun, 2020). One of the metabolites produced during progesterone metabolism is allopregnanolone (ALLO). Both progesterone and allopregnanolone are neurosteroids which can be synthesized within the nervous system or accumulate in the brain via systemic circulation after being derived by the gonads (in females) and by the adrenal glands (in both sexes) (Ghoumari et al., 2014). There is considerable evidence from both clinical studies and animal experiments regarding the effect of progesterone (and allopregnanolone) on human and animal EEG, particularly in exacerbating pathological activity (Grünewald et al., 1992; Budziszewska et al., 1999; van Luijtelaar et al., 2001). Allopregnanolone, a neuroactive steroid, has the ability to modulate neuronal excitability through both genomic (i.e., through regulating gene expression) and nongenomic means (i.e., modifying ion conductance, second messengers, and activating signalling pathways) (Zamora-Sánchez and Camacho-Arroyo, 2022). At the nongenomic level, its effect on GABAa receptors depends on its concentration. At nanomolar concentrations, ALLO is a positive allosteric modulator of the GABAa receptor (Pinna et al., 2000; Puia et al., 2003; Lambert et al., 2009). Specifically, it increases the probability of the channel being in the open state and enhances the receptor’s response to GABA by increasing its e?icacy (Bromfield et al., 2006; Majewska et al., 1986). At micromolar concentrations, ALLO can activate GABAa receptors independently of GABA (Liu et al., 2002; Lambert et al., 2009). Its overall effect is to potentiate GABA action particularly by decreasing the rise time and increasing the decay time of the evoked current, an effect consistently observed across various experimental conditions (Majewska et al., 1986; Paul and Purdy, 1992; Sullivan and Moenter, 2003; Strömberg et al., 2006; Lu et al., 2023). Additionally, under some conditions, an increase in current amplitude is also observed (Paul and Purdy, 1992; Sullivan and Moenter, 2003; Majewska et al., 1986).

In our recent work using a conductance-based thalamocortical model, we found that ALLO’s modulation of GABAa-receptor mediated inhibition had an ameliorating effect on spike-wave discharges (Ahmed and Campbell, 2024). This finding appears to contradict established evidence but may reflect limitations in existing experimental models, which do not capture the naturally remitting course typical of most CAE cases (Grünewald et al., 1992; van Luijtelaar et al., 2001). The thalamocortical model employed in our previous study featured a well-defined thalamic component based on Destexhe’s work, but was limited by its lack of a comprehensive cortical representation (Destexhe et al., 1998a). The cortical cells in this model incorporated only basic currents (*Na*^+^, *K*^+^, and an additional slow-*K*^+^ current in pyramidal cells) and lacked region-specific or laminar organization.

In this study, we develop an enhanced minimal conductance-based thalamocortical model with an improved cortical component. Our revised model incorporates a layered cortical structure featuring region-specific pyramidal cells and deep-layer interneurons from layers 5 and 6. These modifications create a more physiologically relevant thalamocortical network while maintaining computational e?iciency. We use this model to investigate how variations in cortical neuronal composition influence seizure dynamics, with a particular focus on allopregnanolone as a neuromodulatory agent in the resolution of spike-wave discharges. Specifically, we examine how different ratios of intrinsically bursting to regular spiking neurons in layer 5 affect the generation, propagation, and maintenance of spike-wave discharges. Furthermore, we explore the impact of enhanced frontocortical connectivity on these different cortical configurations. Our analysis aims to identify distinct network features that may distinguish CAE networks in which spike-wave discharges are resolved by allopregnanolone from those in which such neurosteroid modulation is ineffective. This computational approach offers a valuable means to explore factors underlying divergent disease trajectories, addressing a key gap in understanding due to the lack of a remitting animal model for CAE.

## 2 Materials and methods

### 2.1 Model description

Our 475-cell thalamocortical network consists of single-compartment neuron models of the following types: layer 5 regular spiking (RS), layer 5 intrinsically bursting (IB), layer 6 non-tufted regular spiking (NRS) pyramidals and a deep low-threshold spiking (LTS) interneuron in the cortex, and thalamic reticular cells (RE) and thalamocortical cells (TC) in the thalamus. The membrane potential of each cell type is described by the following equations:

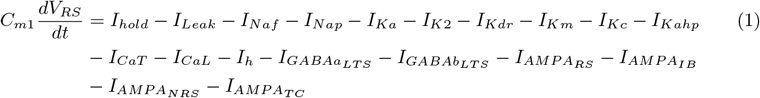

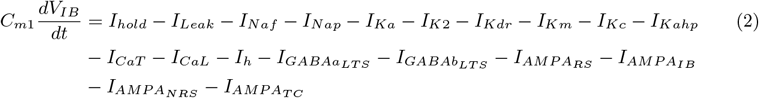

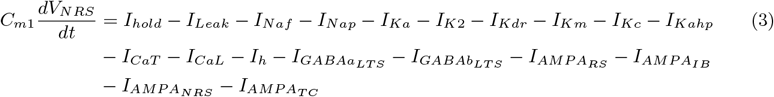

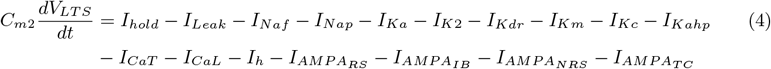

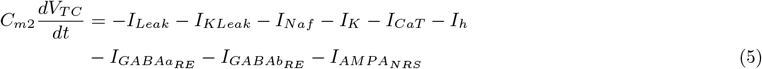

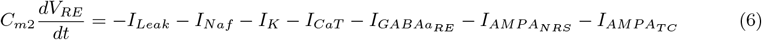

where *V*_*i*_ is the membrane potential for *i* = RS, IB, NRS, LTS, TC, RE, *C*_*m*1_ = 0.9 and *C*_*m*2_ = 1.0 is the specific membrane capacitance given in *µ*F*/*cm^2^. *I*_*hold*_ is the input current required to hold and maintain the voltage of the neuron (also used in this context for setting the resting membrane potential). A consistent set of units were maintained such that voltage is given in mV, ionic currents in mA/cm^2^, membrane conductance densities in S*/*cm^2^, and time in msec. Note that while input currents are given in nA, they are converted to a current density by dividing by the cell compartment’s area.

All cortical cells have the same set of ionic currents with different kinetics in some instances. These consist of both transient and persistent sodium currents, potassium currents of the delayed rectifier, transient, slowly inactivating, and *Ca*^2+^-dependent types, in addition to both lowand high-threshold *Ca*^2+^ currents. The models for these cells were adapted from our previous work (Ahmed, 2019), which modified single-neuron models from Traub et al. (Traub et al., 2005). The original model by Traub et al. featured multi-compartment neurons in a large-scale network, successfully replicating various thalamocortical phenomena. In our work, we simplified these complex neuronal models to single-compartment representations of the soma while preserving essential dynamics. This reduction required careful recalibration of membrane conductance densities to maintain physiologically realistic behaviour. In contrast, the thalamic cells were based on the model by Destexhe et al. (Destexhe et al., 1998a), which describes thalamic neurons with fewer currents but still exhibits bursting behaviour due to the presence of the low-threshold calcium current, with slower kinetics in the RE cells. All intrinsic currents follow the same formalism, described by the following equation:

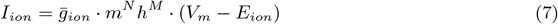

where 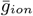 represents the maximal conductance, and *m* and *h* represent the voltage gating of the ion channels. All synaptic currents are described by the following equations:

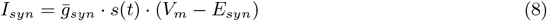

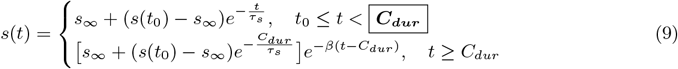

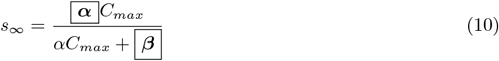

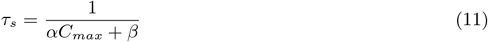

where 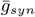 denotes the maximal synaptic conductance, and AMPA/GABAa- and GABAb-mediated synapses are described appropriately by the synaptic conductance function, *s*(*t*) which denotes the fraction of open synaptic channels at time *t*. The time of a presynaptic spike arrival is denoted by *t*_0_, while *α* and *β* denote the forward (binding) and backward (unbinding) rate constants describing the binding of neurotransmitter, respectively. The duration of the neurotransmitter-mediated pulse is given by *C*_*dur*_, and *C*_*max*_ refers to the maximal neurotransmitter concentration value. The details of the individual currents, synaptic dynamics, and model parameters are provided in Appendix A and Appendix B, respectively.

The network consists of the following cell populations: 25 layer 5 RS pyramidals, 75 layer 5 IB pyramidals, 75 layer 6 non-tufted RS pyramidals, 100 deep LTS interneurons, 100 thalamocortical cells, and 100 thalamic reticular cells. At baseline, a higher proportion of bursting-type pyramidal cells compared to regular-spiking cells in layer 5 was used to both introduce asymmetry in Layer 5, as well as model the frontal cortex. All cell layers were of equal size, except for layer 6, which had a reduced population size relative to layer 5 to reflect its typically lower density of pyramidal cells (DeFelipe et al., 2002). Each layer of cells is arranged in one dimension as shown in Figure 1 with synaptic connectivity varying between cell types as described. Cortical input into the thalamus is only from layer 6 of the cortex with excitatory synapses formed with ascending thalamic axons. All excitatory connections in the network are mediated by AMPA receptors, and inhibitory connections are mediated by either GABAa only or a combination of GABAa and GABAb receptors.

**Figure 1.**
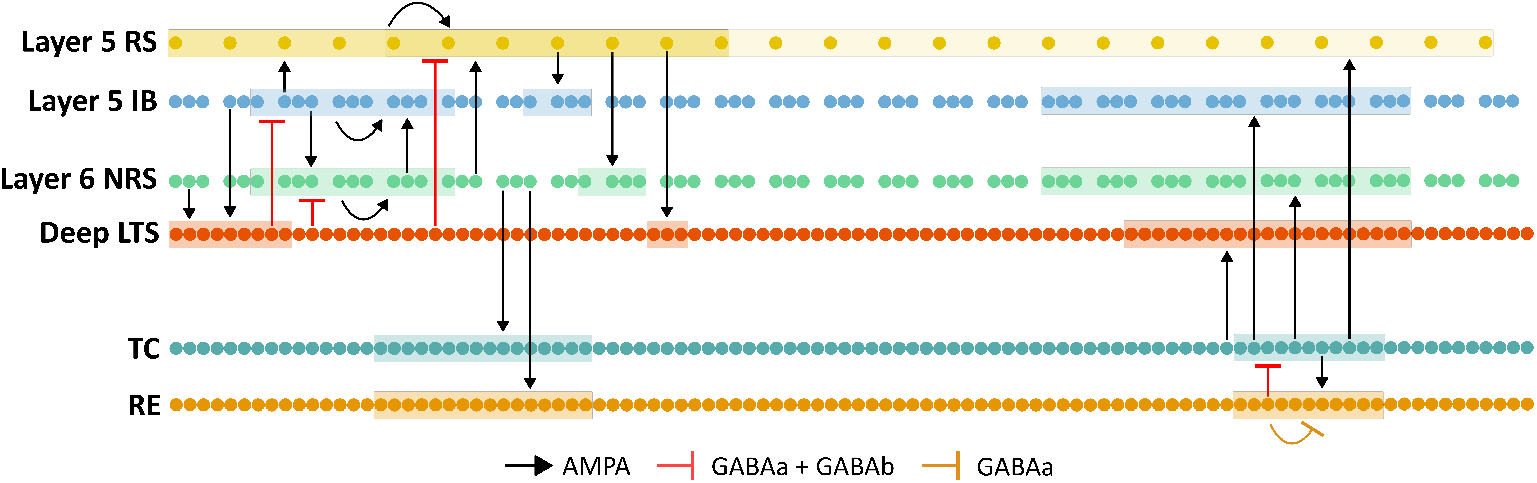
Schematic representation of the thalamocortical network. Each neuron is represented by a dot with synaptic connectivity within and between layers as shown. The number of postsynaptic neurons each neuron connects to varies by layer and is represented by the highlighted groups as shown.

Due to the unequal number of cells in some layers, the number of presynaptic neurons each postsynaptic neuron connects to (*nPrePost*) was layer-dependent, as given in Table 1. In each network layer, presynaptic cells were indexed relative to the indexing of postsynaptic cells. Within the thalamus and the cortex, each neuron connects to postsynaptic neurons within a radius of 11 indices, while between the thalamus and cortex, each presynaptic neuron connects with neurons within a radius of 21 postsynaptic indices. This cell count affects the weight of individual synapses which is defined by 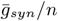 *nP rePost* where 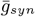 is the maximal synaptic conductance for a particular synapse, as given in Table 2.

**Table 1:** The number of presynaptic neurons that each postsynaptic neuron connects to (*nPrePost*) in a network with the following architecture: 25 layer 5 RS cells, 75 layer 5 IB cells, 75 layer 6 NRS cells, 100 deep layer LTS cells, 100 TC cells, and 100 RE cells. For example, nRSIB = 4 means that each IB cell receives input from 4 RS cells.

| $nPrePost$ | Number of cells | $nPrePost$ | Number of cells |
| --- | --- | --- | --- |
| nRSRS | 11 | nLTSRS | 11 |
| nRSIB | 4 | nLTSIB | 11 |
| nRSNRS | 4 | nLTSNRS | 11 |
| nRSLTS | 3 | nTCRE | 11 |
| nIBIB | 11 | nTCNRS | 21 |
| nIBRS | 11 | nTCIB | 21 |
| nIBNRS | 11 | nTCRS | 21 |
| nIBLTS | 9 | nTCLTS | 21 |
| nNRSNRS | 11 | nRERE | 11 |
| nNRSRS | 11 | nRETC | 11 |
| nNRSIB | 11 |  |  |
| nNRSLTS | 9 |  |  |
| nNRSTC | 16 |  |  |
| nNRSRE | 16 |  |  |

**Table 2:** Maximal synaptic conductance, 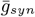 parameters (in *µ*S), for each synapse type, chosen to produce a default network state exhibiting spindle oscillations. Values for GABAa synapses reflect conductance under Control receptor activity.

| Pre-synaptic neuron type | Post-synaptic neuron type |  |  |  |  |  |
| --- | --- | --- | --- | --- | --- | --- |
|  | RS | IB | NRS | LTS | TC | RE |
| RS | 0.3 | 0.7 | 0.3 | 0.1 | - | - |
| IB | 1 | 0.3 | 0.05 | 0.08 | - | - |
| NRS | 0.1 | 0.3 | 2 | 0.2 | 0.02 | 2.4 |
| LTS | 0.09 (GABAa) | 0.1 (GABAa) | 0.75 (GABAa) | - | - | - |
|  | 0.03 (GABAb) | 0.03 (GABAb) | 0.03 (GABAb) |  |  |  |
| TC | 1 | 0.7 | 1.2 | 0.4 | - | 0.2 |
| RE | - | - | - | - | 0.02 (GABAa) | 0.2 |
|  |  |  |  |  | 0.04 (GABAb) |  |

After allowing the network to reach steady state, a 1 nA stimulus was applied for 100 ms to a group of five layer 6 NRS neurons, inducing additional oscillations. Network behaviour and states were characterized using these triggered oscillations. The model fitting parameters, given in Tables 2–4 were selected based on the method described in Section 2.2, and were intended to be interpreted qualitatively. The same set of fitted parameters was used to model both the default and diseased states. In the default mode, the network exhibits spontaneous oscillations driven by spontaneous discharges in TC neurons, while in the diseased mode, the model exhibits synchronized spike-wave discharges due to alterations in the strength of cortical inhibition.

### 2.2 Model fitting procedure

The model fitting procedure aimed to produce target behaviour qualitatively similar to the models upon which we have built this work (Destexhe, 1998; Traub et al., 2005). To assess model fitness, we defined specific evaluation metrics that characterize network behaviour in two critical states: healthy spindles and pathological spike-wave-discharges (SWDs). Our previous model guided the qualitative and quantitative criteria for both states (Ahmed and Campbell, 2024), the details of which are provided in (Ahmed, 2025).

Since the thalamus component was well-defined in (Destexhe, 1998), we kept it unchanged, including its synaptic weights and other related parameters. Similarly, all parameters associated with GABAb synapses were preserved as defined in (Destexhe, 1998). Instead, we focused on fitting parameters corresponding to AMPA and GABAa synapses within the cortex and between the cortex and thalamus. For each synapse, we sought the optimal set of parameters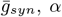, and *β*—as defined in Equations (8)–(10)—that would enable the network to display a default mode of spindle oscillations while maintaining the ability to transition to a spike-wave discharge (SWD) state under cortical disinhibition. This resulted in a 63-dimensional parameter search space. The choice of 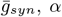, and *β* as fitting parameters was guided by prior optimization and dimensionality reduction efforts. Furthermore, the search space was restricted by narrowing each parameter to select values within particular ranges. Model fitting proceeded in multiple stages, beginning with the cortical layer 6thalamus circuit, then adding cortical layer 5 cells, one population at a time. At each stage, synapses were incrementally added to the network, with the corresponding parameter triplet 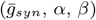 optimized through a grid search to maintain target behaviour before introducing the next synaptic connection. Throughout, the model was periodically evaluated under two conditions—total cortical inhibition and total disinhibition—to ensure emergence of expected behaviours. Model outputs were assessed using an automated procedure with predefined criteria, supplemented by manual inspection of individual neuron firing patterns for physiological plausibility. We tested our model’s robustness by exploring parameter clouds with 30–200% variation around each nominal triplet, with the variation range depending on synapse type and insights from the sequential fitting process. For synapses that strongly influenced network behaviour, we tested smaller neighbourhoods around the nominal values. Within these clouds, multiple triplets met our filtering criteria, indicating local robustness. Further details on the selection of fitting parameters and the staged fitting procedure can be found in (Ahmed, 2025).

### 2.3 Tools for analysis

The model was implemented in NetPyNE, a Python package developed by Dura-Bernal et al. that allows for high-level specification of network connectivity of biological neuronal networks and simulations using the NEURON simulator (Dura-Bernal et al., 2019).

#### 2.3.1 Local field potential and spectral density analysis

The local field potential (LFP) was calculated using a point current source model in which current sources are treated as if they originate from a single point. Due to memory limitations, only a subset of the Layer 5 RS and IB populations was recorded from and used in the LFP calculation. For each population, neurons are arranged in a single line 20 *µ*m apart. The extracellular recording site is considered to be 50 *µ*m opposite to the center of the line, and the LFP at this site was calculated from post-synaptic currents using the following equation:

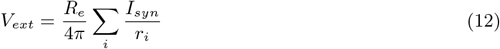

where *V*_*ext*_ is the potential at a defined extracellular site, *R*_*e*_ = 230 *cm is the extracellular resistivity, *I*_*syn*_ is the post-synaptic current, and *r*_*i*_ is the distance between the location of the postsynaptic currents and the extracellular site. The LFP traces were first detrended using the *SciPy* Python library to eliminate very low-frequency components that could contaminate the power spectrum and cause spectral leakage. The traces were then filtered using a Butterworth bandpass filter, with cutoff frequencies set at 1 Hz and 15 Hz. Power spectral density (PSD) analysis was performed on the filtered LFP traces using Welch’s method with a sampling frequency (*fs)* of 10 kHz. A Hann window was applied to segments of 20000 samples, with 1000 samples of overlap between consecutive segments.

### 2.4 Modelling the effect of genetic mutations on GABAa current

Mutations in genes encoding GABAa receptor subunits, including the *GABRG*2 gene, have been implicated in childhood absence epilepsy (Kananura et al., 2002; Kang and Macdonald, 2004; Ito et al., 2005; Marini et al., 2003). Functional studies of mutant receptors, particularly those containing the *γ*2(R82Q) and *γ*2(R43Q) subunits, have demonstrated reduced receptor surface expression in cortical pyramidal neurons and decreased GABAa receptor current amplitudes (Kang and Macdonald, 2004; Macdonald et al., 2010). These alterations are consistent with a reduction in neuronal feedforward inhibition within cortical circuits (Bianchi et al., 2002; Currie et al., 2017). In our model, the effects of the *GABRG*2 mutation were implemented specifically in cortical pyramidal neurons across both layers by modifying the parameter corresponding to the maximum conductance of the GABAa current 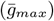,as outlined below:

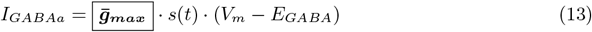

### 2.5 Modelling the effect of allopregnanolone

The effect of allopregnanolone (ALLO) was modelled at the level of synapses, particularly GABAa receptor-mediated synapses, by adjusting the 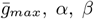,and *C*_*dur*_ parameters, as highlighted in Equations (9)–(11) and Equation (13).

Modifications to the highlighted parameters were informed by fitting our synapse model to experimental data, using the relative change between control GABAa receptor activity and activity after ALLO application. It is important to note that the available experimental data vary considerably in methodology across studies, including differences in the species of central nervous system neurons used, and the concentrations of both GABA (0.005 to 0.1 mM) and ALLO (30 to 1000 nM). While the physiological concentration of GABA released into the synaptic cleft ranges from 1 to 10 mM, the experimental data used to model the effects of ALLO are based on GABA concentrations that are at least an order of magnitude lower (Brickley and Mody, 2012; Overstreet et al., 2002; Tretter and Moss, 2008; Mozrzymas et al., 2003). Similarly, physiological ALLO levels in rat cortex typically range from 1 to 20 nM, yet experimental protocols generally apply concentrations that are one to two orders of magnitude higher (Pinna et al., 2000; Kelley et al., 2011). The lack of significant differences in response amplitude (i.e., relative change in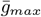) following the application of ALLO in studies using low mM concentrations of GABA could perhaps be explained by these variations. Given these discrepancies, our modelling of these effects is meant to be interpreted qualitatively. The details of the curve fits can be found in Appendix B.1.

While the *C*_*dur*_ parameter corresponding to Control synapses 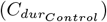 for each GABAa receptor-mediated synapse was set to 0.3 ms, the values of 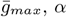, and *β* corresponding to Control synapses (i.e., 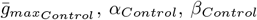) were taken from the values given in Tables 2–4. For post-ALLO synapses, each parameter was modified based on the relative changes between control GABAa receptor activity and activity following the application of ALLO, as follows:

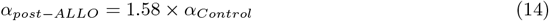

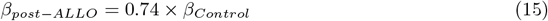

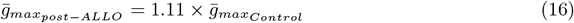

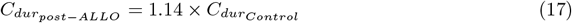

### 2.6 Modelling regional neuronal heterogeneity and enhanced frontocortical connectivity

To model the regional differences in neuronal heterogeneity observed experimentally, as discussed previously, we developed two distinct model configurations. Given that layer 5 intrinsically bursting (IB) neurons are more abundant in the frontal cortex and regular spiking (RS) neurons in the parietal cortex, we modelled the frontal cortex using IB neurons while the parietal cortex was modelled using RS neurons. Accordingly, we varied their proportions in each model such that the 95-5 (nIB:nRS) configuration is representative of a cortex most influenced by the frontal cortex, while the 5-95 (nIB:nRS) configuration reflects a cortical component that is most influenced by the parietal cortex. Given the differences in model architecture, the number of presynaptic neurons each postsynaptic neuron connects to (*nPrePost*) differs between the two model configurations, as presented in Tables B3–B4.

Additionally, for each model configuration, we modelled enhanced connectivity in the frontal cortex by modifying the strength of the following synapses, as illustrated in Figure 2: IBIB, IBNRS, and NRSIB. Specifically, we modified the maximal synaptic conductances as follows: 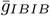 was increased by a factor 7, 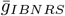 by a factor of 5, and 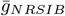 by a factor of 3. A detailed description of the method used to determine these parameter adjustments is provided in Figure B2.

**Figure 2.**
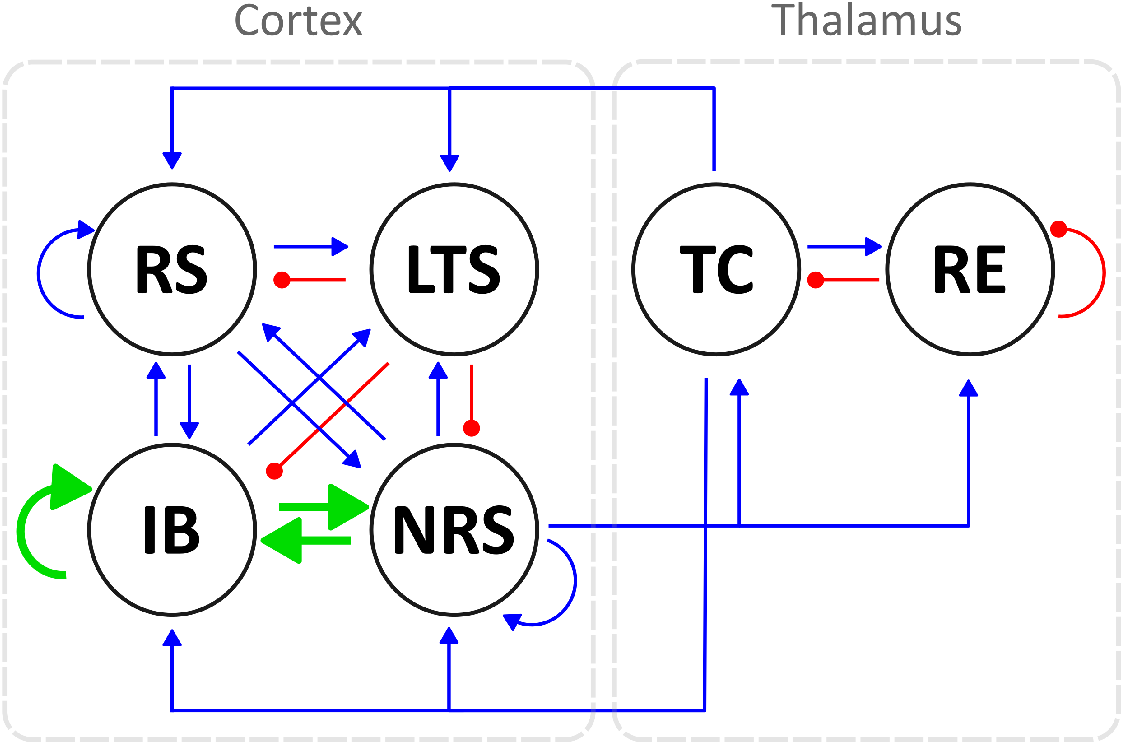
A schematic diagram of the thalamocortical circuit shows connections between cortical neurons of the RS, IB, NRS, and LTS type, and thalamic neurons (TC and RE). Blue arrows indicate excitatory synaptic connections, while red arrows represent inhibitory ones. Enhanced frontocortical connectivity is illustrated with thicker green arrows, which also represent excitatory synapses.

## 3 Results

### 3.1 Baseline state characterization and allopregnanolone effects on the fitted model

The goal of our model fitting was to replicate behaviour that is qualitatively consistent with that of prior established models. Specifically, the model needed to exhibit two critical baseline states: a healthy state characterized by spindle oscillations with a network frequency of 7–10 Hz, and a diseased state marked by spike-wave discharges (SWD) with a network frequency of 2.5–5 Hz. The 75-25 (nIB:nRS) model was simulated using GABAa synapses that were all of the Control type, as described in Table 2. At 100% baseline cortical GABAa conductance, the network produces spindle oscillations with a peak network frequency of 7 Hz, as shown in Figures 3A and 3C. In this healthy state, cell populations exhibit organized, alternating bursting patterns—particularly within the TC population—with moderately synchronous activity across other populations, as shown in Figure 3B. This activity is reflected in the LFP trace as a low-amplitude rhythmic oscillation. The network could be transformed to a diseased state by means of cortical disinhibition. By reducing GABAareceptor mediated inhibition in the cortex to 10% baseline conductance, the network produces a dominant 5 Hz SWD pattern as shown in Figures 3D and 3F. In this diseased state, neural firing becomes more synchronous across all populations, with distinctive spike-wave complexes visible in the LFP, as shown in Figure 3E. The reduction in cortical GABAa conductance was determined by examining how changes in conductance influence network frequency and the relative power in the SWD frequency range, as shown in Figure 4.

**Figure 3.**
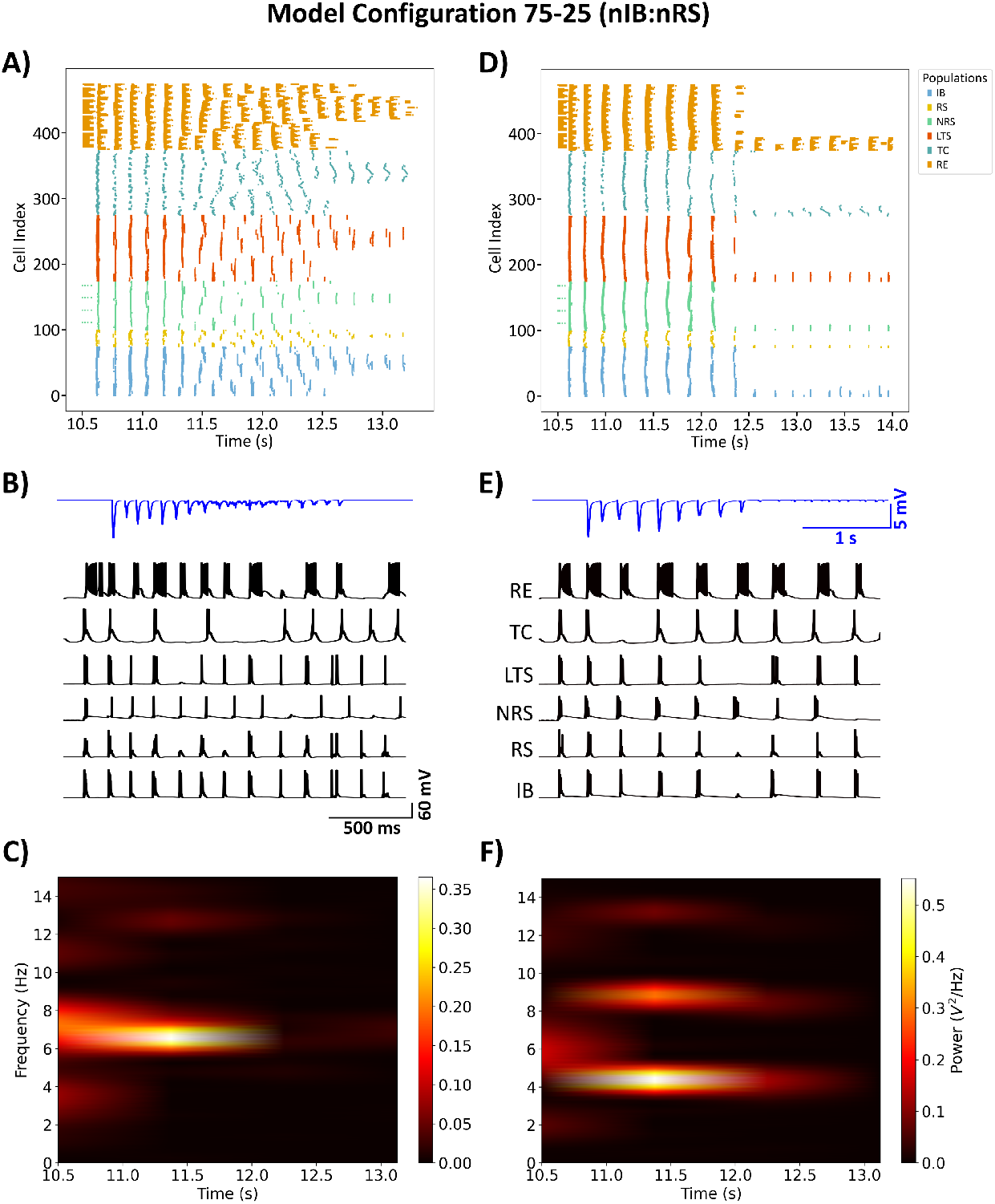
Raster plot of the 72-25 (nIB:nRS) fitted model demonstrates spindle oscillations (A), transitioning to spike-wave discharges (D) through cortical disinhibition, modelled as a reduction in baseline GABAa conductance from 100% to 10%. (B) and (E): Sample voltage traces from cell index 0 in each population, with LFP traces (in blue) computed using the method outlined in Section 2.3. The network activity during spindle oscillations exhibits a peak power at 7 Hz (C), whereas the SWD state shows peak power at 5 Hz (F). All simulations were conducted using Control GABAa-receptor mediated synapses.

**Figure 4.**
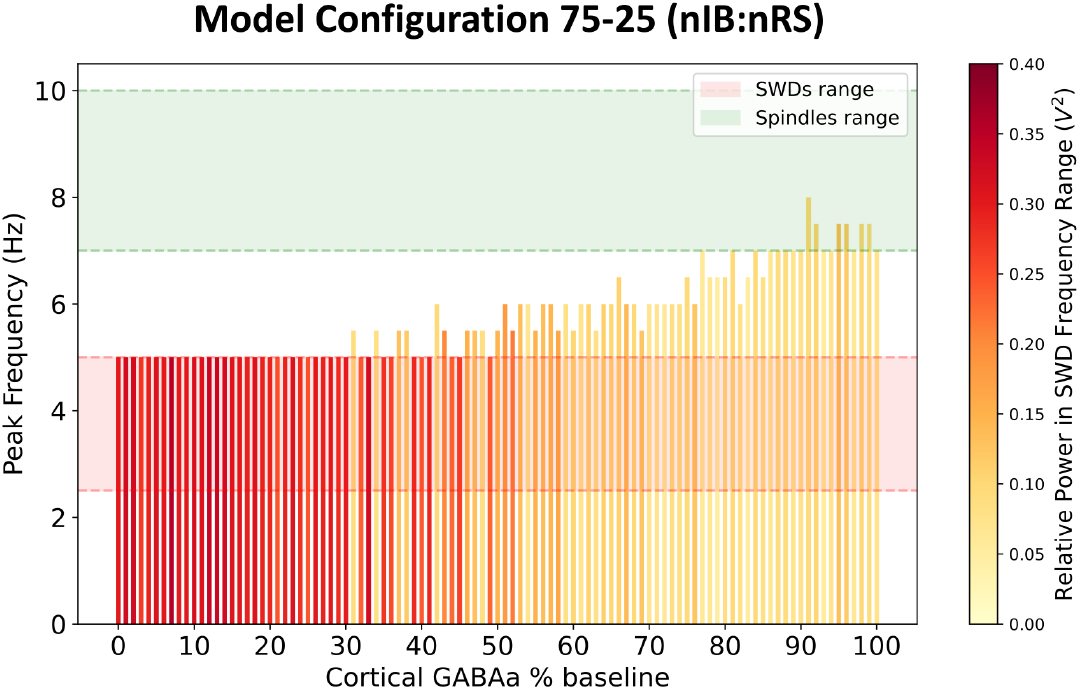
Network frequency response to varying levels of cortical GABAa-receptor mediated inhibition. The colour of each bar represents the relative power within the frequency range associated with SWDs.

Next, we modelled the effect of allopregnanolone (ALLO) by altering parameters associated with the GABAa current described in Section 2.5. The network was initialized to exhibit SWDs, as in Figures 3D–F, with cortical GABAa conductance reduced to 10% of baseline. The effect of ALLO was implemented uniformly across all GABAa synapses in the model. Our results show that allopregnanolone effectively suppresses pathological SWDs by altering the temporal firing patterns of neural populations throughout the thalamocortical circuit. As shown in Figure 5, there is a shift in the dominant network frequency to 7 Hz, which falls within the frequency band associated with normal spindle oscillations. This transition is also characterized by alternating bursts of synchrony across populations which is reflected in the low-amplitude filtered LFP trace. The shift from pathological to physiological dynamics is consistent with our previous findings and supports the therapeutic potential of positive GABAa modulators in absence epilepsy.

**Figure 5.**
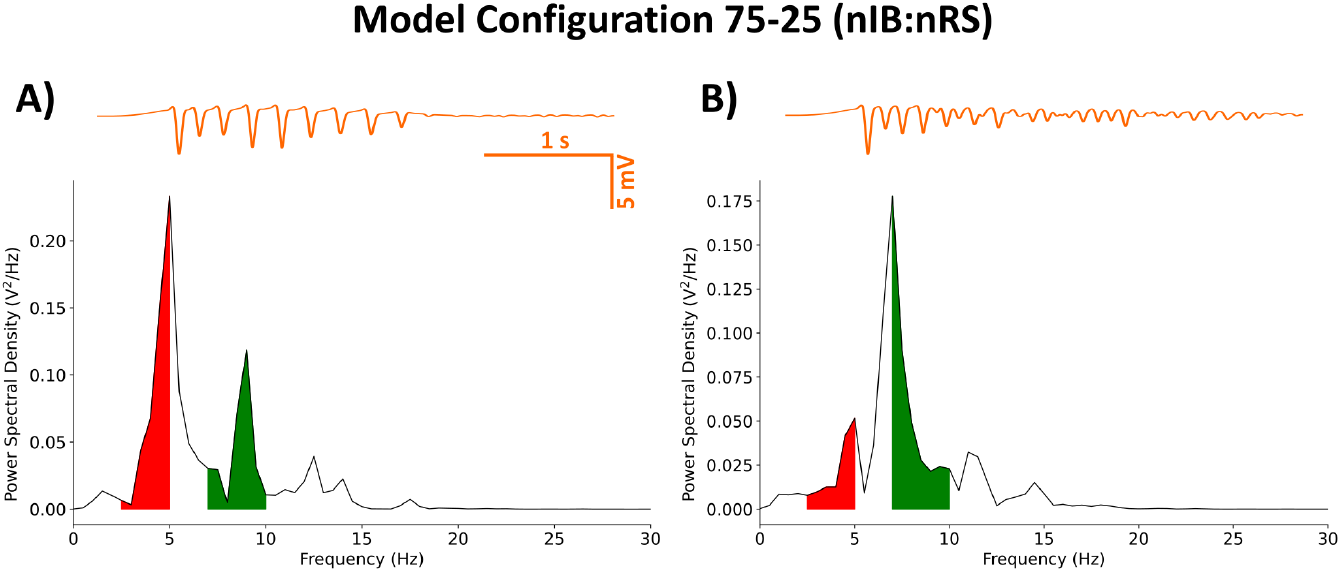
Power spectral density of the 75-25 (nIB:nRS) fitted model using Control synapses (A) and post-ALLO synapses (B), calculated from filtered LFP traces (in orange), as described in Section 2.3. The network was initialized with cortical disinhibition (10% baseline cortical GABAa conductance). Shaded regions indicate relative power within the frequency ranges associated with SWDs (2.5–5 Hz; red) or Spindle oscillations (7–10 Hz; green).

### 3.2 Regional neuronal heterogeneity in the cortex does not alter ALLO-mediated network dynamics

Using the previously described frontal cortex composition (95-5 nIB:nRS) and parietal cortex composition (5-95 nIB:nRS) in Section 2.6, we next examined whether regional neuronal heterogeneity affects the network’s propensity to generate the two critical network states of interest—healthy spin-dles and pathological SWDs. Additionally, we investigated whether the effect of ALLO differs across networks with different cortical compositions, and importantly, whether one network composition is more prone to non-resolution than the other.

We first investigated the excitability profiles of both network configurations by examining network frequency responses to varying levels of GABAa-receptor mediated inhibition in the cortex, as shown in Figure 6. Both models display a transition from SWD to spindle activity as cortical inhibition increases. However, the thresholds at which this transition occurs differs slightly between the two, with the 95-5 (nIB:nRS) model exhibiting more intermediate oscillatory states compared to the 5-95 (nIB:nRS) model. The latter surprisingly maintains SWD activity across a broader inhibition range and requires stronger inhibition to shift to spindle-generating states. To further explore the influence of cortical composition, we simulated both networks under the same conditions as in Section 3.1— 100% and 10% baseline cortical GABAa conductance—to compare their abilities to sustain either physiological spindles or pathological SWDs, and to determine whether baseline dynamics differ across the two models. As shown in Figure 7, both network configurations exhibit qualitatively similar spectral characteristics, across the default mode and under cortical disinhibition. The spectral power distributions show similar frequency bands of activity (both exhibiting peak network frequencies of 7.5 Hz and 5 Hz under each condition), with only subtle differences in power distribution, likely due to variations in oscillatory patterns, as reflected in the LFP traces. This is also characterized by prominent power spectral density peaks within the 7–10 Hz (Figures 8A1 and 8C1) and 2.5–5 Hz ranges (Figures 8A2 and 8C2). At 100% baseline cortical GABAa conductance, both models exhibit 7.5 Hz peak network frequencies with similar relative power in the spindles frequency range (Figure 8B1). Similarly, at 10% baseline conductance, both models exhibit network frequencies peaking at 5 Hz with similar relative power in the SWDs frequency range (Figure 8B2). These results suggest that despite substantial variation in cortical neuronal composition between models, the fundamental network dynamics remain conserved. This suggests that our fitted model (75-25 nIB:nRS configuration) is robust to changes in network architecture in producing baseline network states.

**Figure 6.**
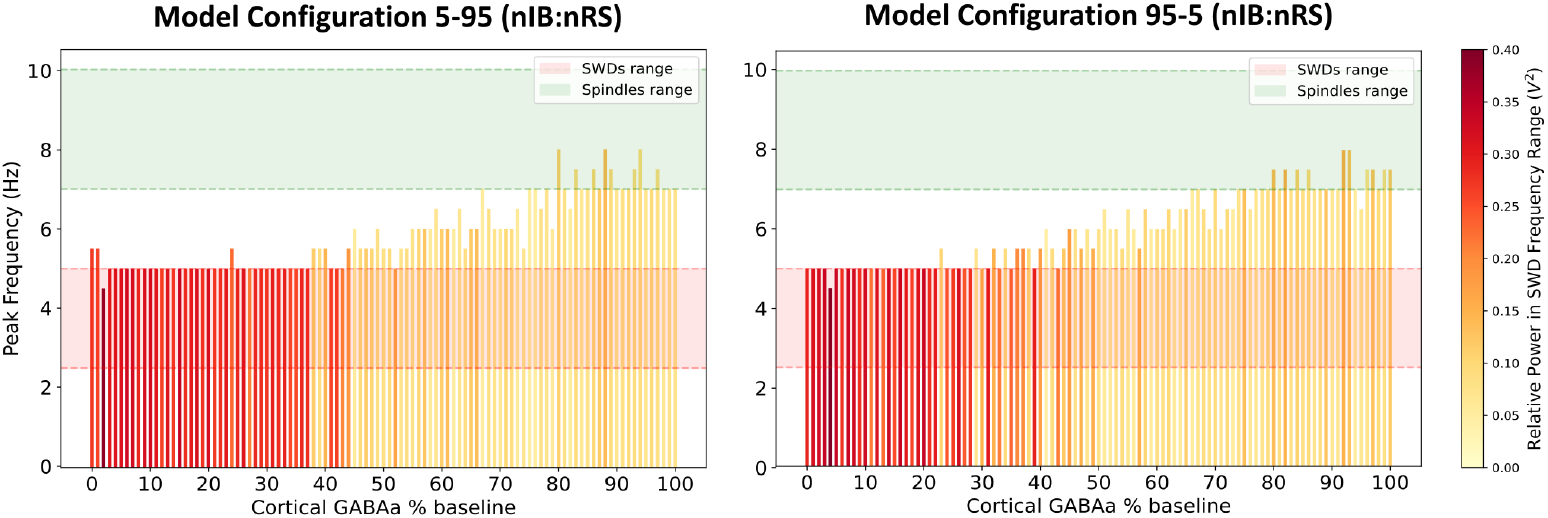
Network frequency response to varying levels of cortical GABAa-receptor mediated inhibition, using the 5-95 (left) and 95-5 (right) (nIB:nRS) model configurations. The colour of each bar represents the relative power within the frequency range associated with SWDs.

**Figure 7.**
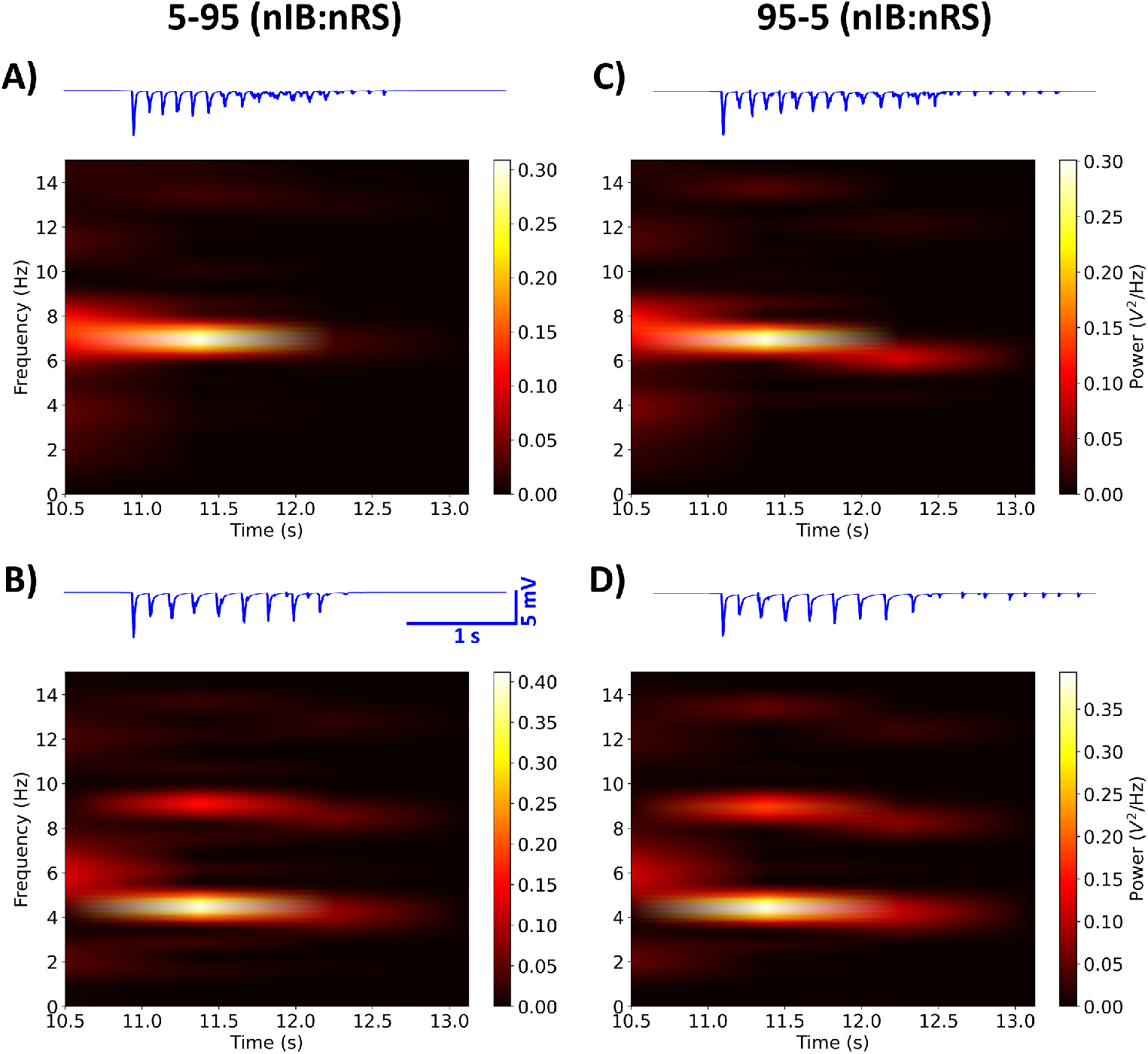
Spectral power analysis showing time-frequency representations of network activity for the 5-95 and 95-5 (nIB:nRS) model configurations, using 100% (A, C) and 10% (B, D) baseline cortical GABAa conductance. For both models, fully inhibited networks exhibit maximal power at 7.5 Hz, while partially disinhibited networks show maximal power at 5 Hz. LFP traces (in blue) were calculated using the population-based method described in Section 2.3.

**Figure 8.**
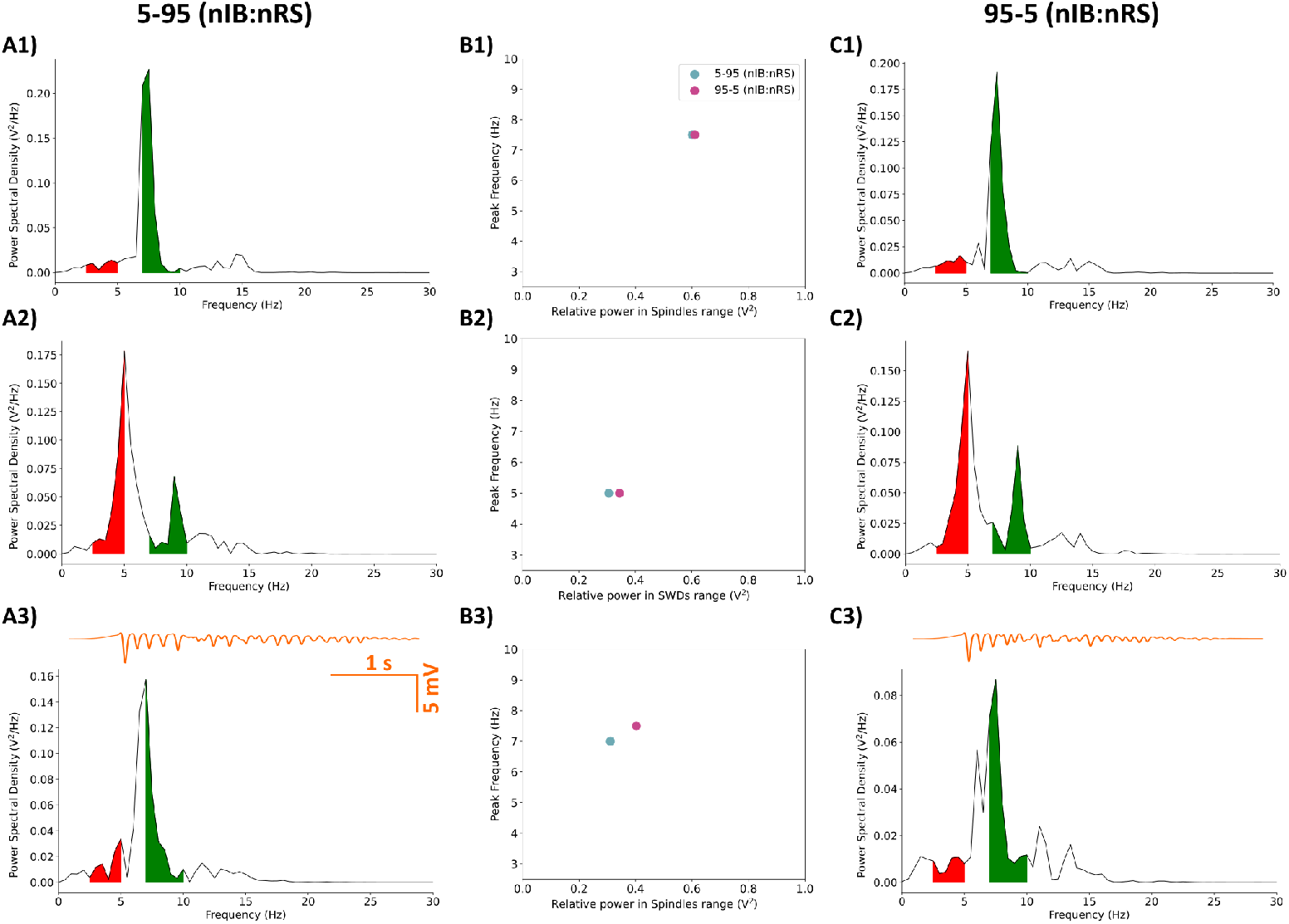
Power spectral density of the 5-95 (A1–A3) and 95-5 (C1–C3) (nIB:nRS) model configurations, calculated from filtered LFP traces (in orange). Shaded regions indicate relative power within the frequency ranges associated with SWDs (2.5–5 Hz; red) or Spindle oscillations (7–10 Hz; green). (A1, C1): Networks with Control synapses and full cortical inhibition, exhibiting spindle oscillations at 7.5 Hz. (A2, C2): Networks with Control synapses and reduced cortical GABAa conductance (10% baseline), exhibiting SWDs at 5 Hz. (A3, C3): Networks initialized with cortical disinhibition as in (A2, C2) but using post-ALLO synapses, showing transition to a physiological state under the effect of allopregnanolone. (B1): Relationship between relative power in spindles range and peak network frequency for fully inhibited networks. (B2): Relationship between relative power in SWDs range and peak network frequency for partially disinhibited networks. (B3): Relationship between relative power in spindles range and peak network frequency for networks under the effect of ALLO.

Following the approach in Section 2.5, we modelled the effect of ALLO in all GABAa synapses, in each model configuration by first initializing to the diseased state. After implementing post-ALLO synapses in each compromised network, we observe a restorative effect, with each network transitioning back to spindle oscillations similar to the control state, as shown in Figures 8A3 and 8C3. This transition is characterized by peak network frequencies of 7 Hz and 7.5 Hz for the 5-95 and 95-5 (nIB:nRS) models respectively, with comparable relative power in the spindles frequency range (Figure 8B3).

While the differences between the two model configurations is small, there is a more obvious difference between the control network states with cortical GABAa at 100% and network states under the effect of ALLO. Specifically, the relative power in the spindle frequency range is higher in the control condition as compared to the post-ALLO state (Figures 8B1 and 8B3). This is mainly due to the more sharply defined spectral density peak curves in the control condition, resulting in greater area under the curve within the spindle frequency range. In the post-ALLO state, the spectral profiles appear more spread out, resulting in relatively less concentrated power contribution falling within the spindles frequency range, despite restoring the dominant network frequency. Overall, these results suggest that, despite differences in the composition of cortical cell types (with one network having a higher proportion of IB cells), both networks respond similarly to cortical GABAa modulation and the effect of allopregnanolone, with neither showing significantly greater vulnerability to the non-resolution of spike-wave discharges.

### 3.3 Enhanced frontocortical connectivity alters SWD resolution based on network architecture

In this part of the study, we focused on modelling the connectivity differences observed in treatment non-responders prior to treatment in newly diagnosed CAE patients, particularly the increased connectivity in the frontal cortex. By simulating a 50-50 (nIB:nRS) model configuration, we aimed to investigate how, in a network with a balanced composition of cortical cell types in layer 5, increased frontocortical connectivity might influence the modulation of SWDs—particularly in the non-resolution of SWDs (i.e., a lack of effect from ALLO on SWDs). Specifically, we modelled enhanced frontal cortical connectivity by modifying the following synaptic conductance parameters with a multiplicative factor to increase synaptic strength: (i)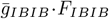; (ii)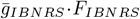; (iii)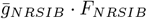. The factors used and the rationale for their selection are detailed in Section 2.6. Additionally, we explored how these enhanced synapses affect the other two model configurations (5-95 and 95-5 (nIB:nRS)) to assess whether, in addition to connectivity differences, the composition of cell types within the network influences its responsiveness to the effects of allopregnanolone.

Building on the approach in Section 3.1, we simulated the 50-50 (nIB:nRS) model with enhanced synaptic strengths, using (*F*_*IBIB*_, *F*_*IBNRS*_, *F*_*NRSIB*_) = (7, 5, 3). We performed the simulation under the two conditions of cortical inhibition (100% and 10% baseline conductance) to investigate the impact of these changes on network activity. As shown in Figure 9, we observed distinct changes in spectral power across different frequencies, depending on the level of cortical inhibition and the presence of allopregnanolone. Specifically, under full cortical inhibition, the network exhibits a maximal power at 5.5 Hz (Figure 9A), marking the first instance where the peak network frequency did not coincide with the frequency exhibiting the highest power. However, the relative power within the spindles range (0.282 V^2^) was greater than that within the range associated with SWDs (0.195 V^2^). When partial disinhibition was applied, the network shifted its maximal power to 4 Hz (Figure 9B). After introducing post-ALLO synapses, the network showed a lack of a restorative effect from ALLO, with the maximal frequency dropping to 2.5 Hz (Figure 9C). The continued SWD-like behaviour is further reflected in the LFP traces of network activity post-ALLO application, especially when compared to network activity under cortical disinhibition, as shown in Figures 9B, C (in blue). Next, we examined the impact of enhanced synaptic activity, with (*F*_*IBIB*_, *F*_*IBNRS*_, *F*_*NRSIB*_) =(7, 5, 3), on the other two model configurations, 5-95 and 95-5 (nIB:nRS). As illustrated in Figure 10A1, B1, C1, both models showed similar network frequencies (6.5 Hz for 5-95 and 7 Hz for 95-5 (nIB:nRS)) and spindle-range relative power (0.339 V^2^ and 0.376 V^2^, respectively) under the condition of 100% baseline cortical GABAa conductance. Reducing cortical GABAa conductance to 10% resulted in comparable network behaviour across models, with peak power at 4 Hz and 3 Hz, respectively, as shown in Figures 10A2, B2, C2. Both of these findings align with results in Section 3.2, particularly the shift from spindle oscillations to SWDs under reduced cortical inhibition. Notably, the 95-5 (nIB:nRS) model consistently exhibited higher relative power across both frequency ranges, regardless of frontocortical connectivity alterations, as illustrated in Figures 11A and 11C. Without enhanced synaptic activity, network frequencies were identical between models and changed similarly across levels of cortical inhibition, as shown in Figures 11B and 11D (orange bars). However, when frontocortical connectivity was enhanced, the 95-5 (nIB:nRS) model demonstrated higher frequency under full inhibition and a lower frequency under partial disinhibition, as illustrated in Figures 11B and 11D (purple bars).

**Figure 9.**
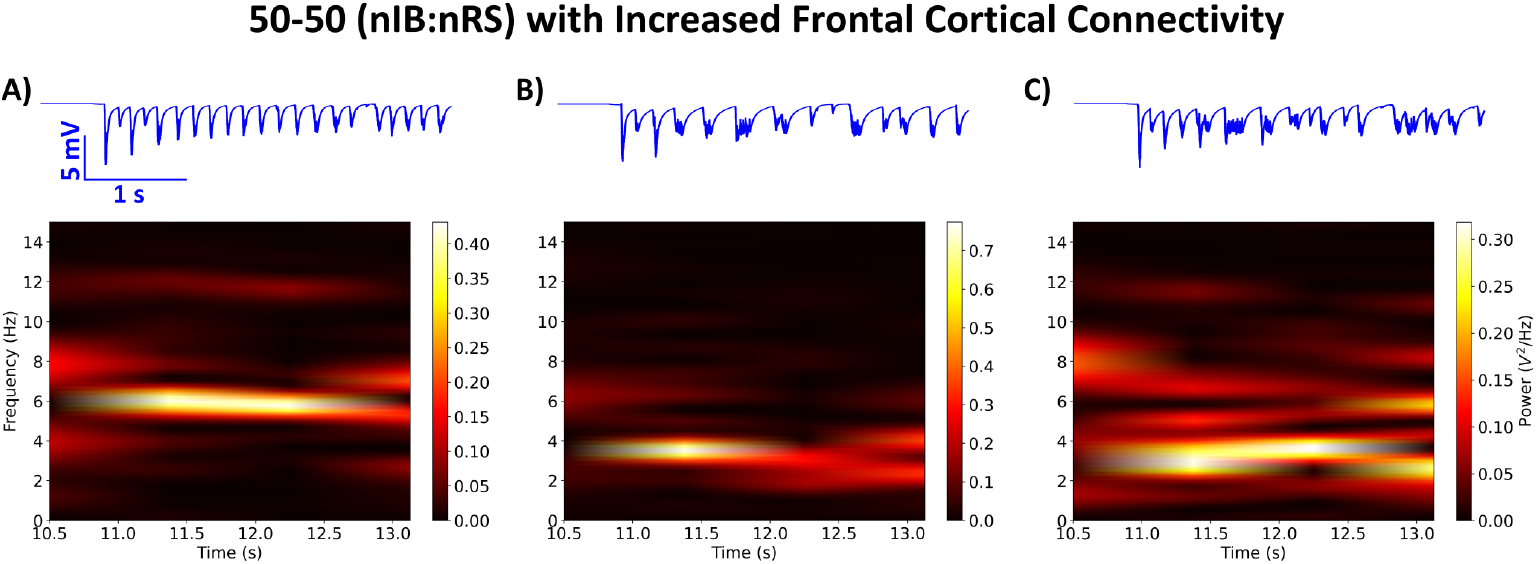
Spectral power analysis showing time-frequency representations of network activity for the 50-50 (nIB:nRS) model configuration with enhanced frontal cortical connectivity, using (*F*_*IBIB*_, *F*_*IBNRS*_, *F*_*NRSIB*_) = (7, 5, 3). (A): Network behaviour using Control synapses and full cortical inhibition exhibits maximal power at 5.5 Hz. (B): Network behaviour using Control synapses under partial disinhibition (10% baseline cortical GABAa conductance) shows maximal power at 4 Hz. (C) Network initialized with cortical disinhibition as in (B) but using post-ALLO synapses shows a lack of restorative effect from ALLO, with a maximal frequency of 2.5 Hz. LFP traces (in blue) were calculated using the method described in Section 2.3.

**Figure 10.**
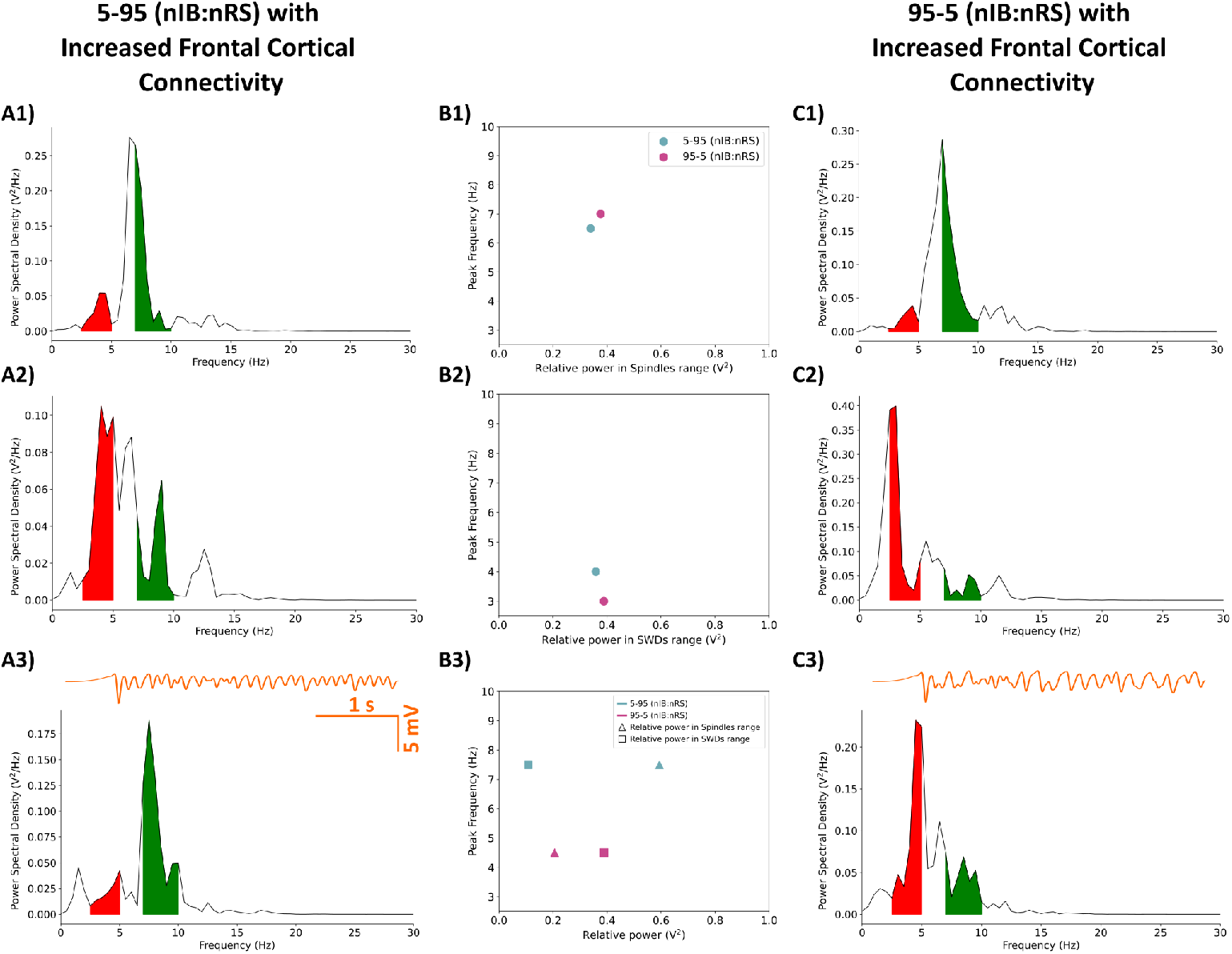
Power spectral density of the 5-95 (A1–A3) and 95-5 (C1–C3) (nIB:nRS) model configurations with enhanced frontal cortical connectivity, calculated from filtered LFP traces (in orange). Shaded regions represent relative power within frequency ranges associated with SWDs (2.5–5 Hz; red) or Spindle oscillations (7–10 Hz; green). (A1, C1): Networks with Control synapses and full cortical inhibition, exhibiting spindle oscillations at 6.5 Hz and 7 Hz for the 5-95 and 95-5 models, respectively. (A2, C2): Networks with Control synapses and reduced cortical GABAa conductance (10% baseline), exhibiting SWDs at 4 Hz and 3 Hz for the 5-95 and 95-5 model, respectively. (A3, C3): Networks initialized with cortical disinhibition as in (A2, C2) but using post-ALLO synapses. The transition to a physiological state under the effect of allopregnanolone occurs for the 5-95 model configuration, but not the 95-5 model. (B1): Relationship between relative power in spindles range and peak network frequency for fully inhibited networks (B1) and networks under the effect of ALLO (B3). (B2): Relationship between relative power in SWDs range and peak network frequency for partially disinhibited networks.

**Figure 11.**
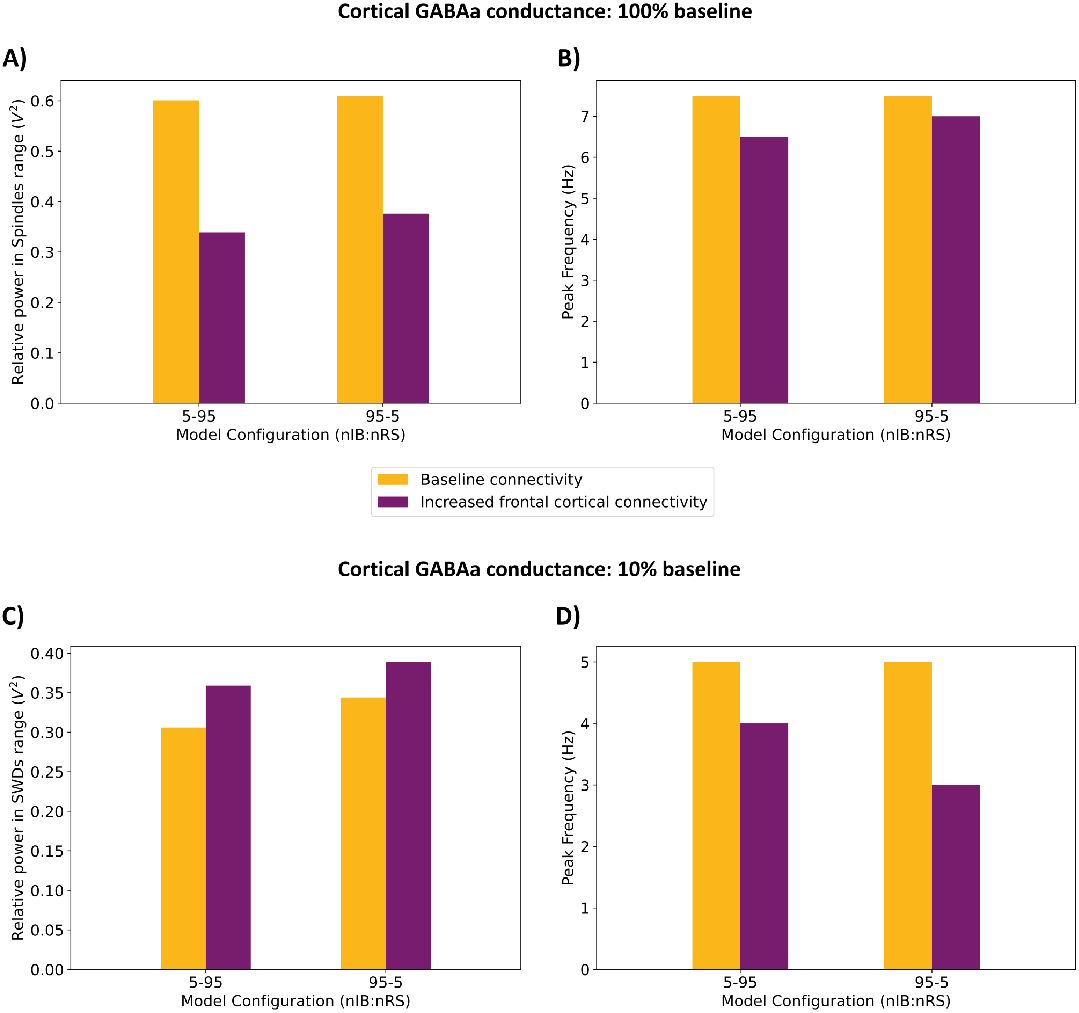
Comparison of network responses between the 5-95 and 95-5 (nIB:nRS) model configurations under two cortical GABAa conductance conditions (100% and 10% baseline). (A, C): Relative power in the spindle/SWDs frequency range. (B, D): Corresponding peak frequencies of network activity. Each condition is shown under baseline (orange) and increased frontocortical connectivity (purple).

Introducing post-ALLO synapses in networks initialized to a pathological state, however, leads to interesting differences between the models. Surprisingly, the effect of ALLO on the 5-95 model mirrors the results from Section 3.2, where allopregnanolone exerts a restorative effect, shifting the network’s maximal power into the spindles range with a peak network frequency of 7.5 Hz, as shown in Figure 10A3. This shift occurs despite there being enhanced frontal cortical connectivity. In contrast, the effect of ALLO on the 95-5 (nIB:nRS) model resembles that seen in the 50-50 (nIB:nRS) model, with maximal power in the SWD-associated frequency range and a peak network frequency of 4.5 Hz.

## 4 Discussion

In this study, we developed a simplified thalamocortical model with a layered cortical structure to investigate how variations in frontocortical connectivity might influence the effectiveness of allopregnanolone in resolving spike-wave discharges in CAE. We modelled the cortical regions with specific neuron firing patterns, using intrinsically bursting (IB) neurons to represent the frontal cortex and regular spiking (RS) neurons to represent the parietal cortex. By exploring two cortical compositions, 5-95 (nIB:nRS) and 95-5 (nIB:nRS), we examined circuits with differing contributions from the frontal and parietal regions. This model allowed us to investigate how neurosteroid modulation, specifically allopregnanolone, interacts with varying cortical compositions to influence seizure dynamics and treatment outcomes.

Our results demonstrate that both physiological and pathological oscillations are maintained by the thalamocortical circuit regardless of which model configuration is used (5-95 or 95-5 (nIB:nRS) model). Under different conditions of cortical inhibition, both models exhibited similar network behaviour, promoting spindles and spike-wave discharge (SWD) activity. This finding aligns with the classification of absence seizures as a generalized epilepsy type, where activity that may originate in one region, such as the parietal cortex, can recruit other cortical areas to sustain seizure activity (Meeren et al., 2002; Polack et al., 2007). Similarly, sleep spindles are known to vary in frequency and cortical source across different brain regions (Purcell et al., 2017). The most striking finding from our study came when we introduced allopregnanolone (ALLO) under conditions of increased frontocortical connectivity. ALLO, a neurosteroid that potentiates GABAa-receptor function, gradually increases during puberty (Fadalti et al., 1999), a period that coincides with the remission of CAE in many cases (van Luijtelaar et al., 2001). In our simulations, both models showed a restorative effect of ALLO, helping them recover from the SWD state at baseline connectivity. However, only the 5-95 (nIB:nRS) model was able to recover when frontocortical connectivity was enhanced. This differential response to ALLO is particularly notable, considering that both models initially display similar behaviour under physiological and pathological states. This suggests that individuals with different neuronal composition profiles may follow distinct clinical trajectories despite similar initial presentations. Specifically, our results suggest that patients with connectivity profiles resembling the 5-95 network (i.e., parietal-dominant configuration) may experience remission following hormonal changes involving allopregnanolone, whereas those with profiles resembling the 95-5 network (i.e., frontal-dominant configuration) may be predisposed to non-remission.

Our results suggest that non-resolving CAE may result not only from increased strength of connections in the frontal cortex but also from the composition of cell types within the network, with a higher proportion of bursting-type cells preventing the therapeutic effects of allopregnanolone. This highlights the significance of individual neuronal components in network dynamics—much like the concept of ictogenic neurons, where even in a network with largely non-pathological connectivity, specific neuronal populations can contribute to pathological activity patterns (Depaulis and Charpier, 2018; Polack et al., 2007). Our findings are consistent with clinical studies using functional connectivity analysis that have identified pre-treatment ictal connectivity differences between patients who ultimately experience remission and those who do not. In particular, increased frontal cortical connectivity has been observed in patients with non-remitting seizures, and treatment nonresponders (Tenney et al., 2018). It is important to note that these clinical studies employ functional connectivity analyses, whereas our modelling involves structural connectivity alterations. Functional connectivity measures the relationship between neural activity in different brain regions, without directly reflecting physical connections (Straathof et al., 2019). On the contrary, structural connec-tivity in our model refers to the actual synaptic connections between neurons. Despite this difference in approach, our ability to reproduce similar results by modifying synaptic strengths suggests that structural alterations may underlie the functional connectivity patterns observed clinically. This provides a potential mechanistic explanation for the clinical heterogeneity observed in CAE outcomes. Some of the limitations of this work include our simplified representation of frontal and parietal cortices, with neuronal firing types (bursting versus regular spiking) serving as the primary distinguishing feature between cortical regions. In reality, both firing types exist across cortical regions, albeit in different proportions (Chagnac-Amitai et al., 1990; Yang et al., 1996). Future models could be improved by incorporating additional distinguishing features of these regions, such as regionspecific distributions of receptors mediating excitation or inhibition. To address the multifactorial nature of this disease, our future directions include exploring whether connectivity differences that prevent SWD resolution hold for other genetic mechanisms contributing to the initial diseased state, beyond the reduced cortical inhibition caused by *GABRG*2 mutations. It would be valuable to explore whether certain genetic mechanisms might better compensate for increased frontocortical connectivity to promote the resolution of SWD activity. Our model is particularly well-suited for investigating differential drug responses based on connectivity profiles. Future studies could test whether specific pharmacological interventions are better suited for particular connectivity patterns. This approach would be similar to existing computational studies that evaluate treatment responses in models with specific genetic mechanisms underlying the disease state, providing insights into potential treatment response trajectories and personalized therapeutic strategies (Knox et al., 2018).

## 5 Conclusion

In conclusion, our study highlights that both neuronal composition and thalamocortical connectivity patterns influence neurosteroid-mediated remission in childhood absence epilepsy. Specifically, our model demonstrates that increased frontocortical connectivity may prevent allopregnanolone-mediated recovery from spike-wave discharge states in networks with higher proportions of intrinsically bursting neurons in deep cortical layers. This may help explain why some patients experience non-remitting seizures despite the hormonal changes associated with puberty. One of our main contributions is that our model provides a computational framework for developing more targeted therapeutic approaches, particularly for patients with distinct excitability and connectivity profiles that may predict a predisposition to poor treatment response or a disease course leading to nonresolution.

## Author Contributions

Each author has substantially contributed to conducting this study and in drafting this manuscript.

## Model availability

Upon publication, the model code will be made publicly available on ModelDB.

## Appendix

### A Model components

All of the neurons were described using a single compartment representing the soma. The soma was modelled as a cylinder with a surface area of *πdl*, where *d* is the diameter and *l* is the length. The diameter was 18 *µ*m for both IB and RS neurons, 16 *µ*m for NRS and LTS neurons, 96 *µ*m for TC neurons, and 70 *µ*m for all RE neurons. The length was 25 *µ*m for both IB and RS cells, 20 *µ*m for NRS and LTS cells, 96 *µ*m for TC cells, and 64.86 *µ*m for RE neurons. The axial resistance, *Ra* was equal to 250 *cm for all cortical cells and 100 *cm for both thalamic neurons. The membrane conductance density distributions as well as the type of current differed between neuron type. Ionic currents were described with the general form given by the following equation:

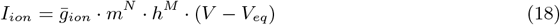

where 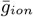 represents the maximal conductance for a particular ion channel, *m* and *h* are gating variables, *N* and *M* are integer powers, and *V*_*eq*_ represents the equilibrium potential. The dynamics of the gating variables were described by one of two general forms: rise and decay rate functions, or a Boltzmann function combined with a voltage-dependent time constant function.

Using rise and decay rate functions, the dynamics were described by the following form:

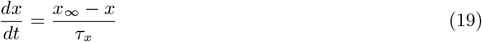

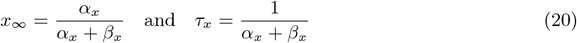

where *x*_*\**_ is the steady-state gating variable function, *τ*_*x*_ represents the time course for approaching the steady state, *α*_*x*_ and *β*_*x*_ represent the forward and backward rate functions.

Alternatively, the gating variable dynamics were described by the following Boltzmann function:

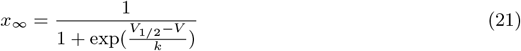

where *x*_*\**_ is the steady-state gating variable function, *V* is the membrane potential, *V*_1*/*2_ is the voltage at half-activation, and *k* is the slope factor representing the steepness of the voltage-dependent transition between closed and open states of the channel. The voltage-dependent time constant function (*τ*_*x*_) describes how quickly the gating variable approaches steady state and its form varies according to the channel type.

The incorporated *Na*^+^, *K*^+^, and *Ca*^2+^ currents included in the model are summarized in Table A1.

The equilibrium potentials were defined as follows: *V*_*Na*_ = 50 mV (all cells), *V*_*K*_ = *\**100 mV (for LTS, TC, RE) and -95 mV (for RS, IB, NRS), *V*_*h*_ = *\**40 mV (for LTS, TC, RE) and -43 mV (for RS, IB, NRS), *V*_*Leak*_ = *\**70 mV (for RS, IB, NRS, TC), -65 mV for LTS and -90 mV for RE neurons, *V*_*Ca*_ was given by intracellular calcium dynamics described in Section A.2.1.

**Table A1:**
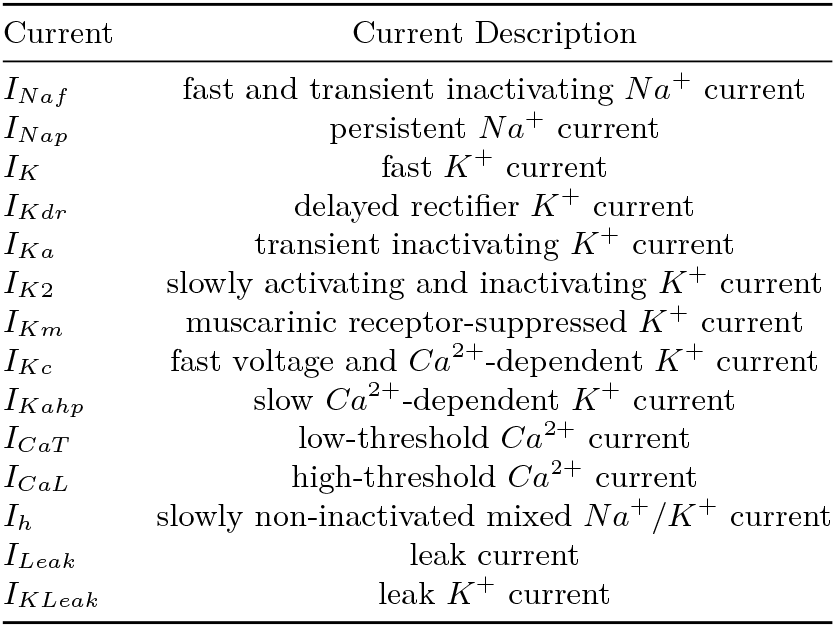
Description of the currents used for all neuron types.

| Current | Current Description |
| --- | --- |
| $I_{Na f}$ | fast and transient inactivating $Na^+$ current |
| $I_{Na p}$ | persistent $Na^+$ current |
| $I_K$ | fast $K^+$ current |
| $I_{K dr}$ | delayed rectifier $K^+$ current |
| $I_{K a}$ | transient inactivating $K^+$ current |
| $I_{K 2}$ | slowly activating and inactivating $K^+$ current |
| $I_{K m}$ | muscarinic receptor-suppressed $K^+$ current |
| $I_{K c}$ | fast voltage and $Ca^{2+}$-dependent $K^+$ current |
| $I_{K ahp}$ | slow $Ca^{2+}$-dependent $K^+$ current |
| $I_{Ca T}$ | low-threshold $Ca^{2+}$ current |
| $I_{Ca L}$ | high-threshold $Ca^{2+}$ current |
| $I_h$ | slowly non-inactivated mixed $Na^+/K^+$ current |
| $I_{Leak}$ | leak current |
| $I_{K Leak}$ | leak $K^+$ current |

#### A.1 Summary of currents

##### A.1.1 I_Leak_

The leak current for all neuron types was described as a simple ohmic current:

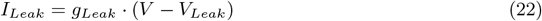

where *g*_*Leak*_ is the constant conductance.

##### A.1.2 I_KLeak_

The leak *K*^+^ current in TC neurons was described by the following Ohmic equation:

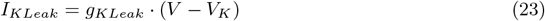

where *g*_*KLeak*_ is the maximal conductance.

##### A.1.3 I_Naf_

The fast *Na*^+^ current was described using a Hodgkin-Huxley style equation:

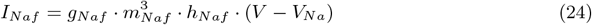

where *g*_*Naf*_ is the maximal conductance, *m*_*Naf*_ is the activation variable, and *h*_*Naf*_ is the inactivation variable.

In neurons of the RS, IB and NRS type, kinetics were based on dynamics described in Traub et al. (Traub et al., 2005):

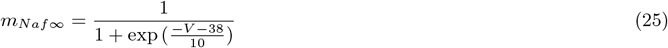

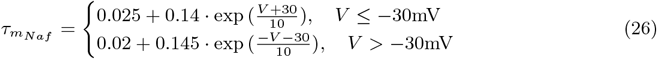

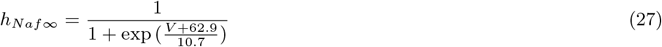

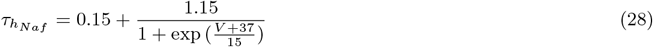

In LTS neurons, the steady state activation function, *m*_*Naf*_, was the same as described above other cortical neurons, while the remaining kinetics were based on dynamics described in Martina and Jonas (Martina and Jonas, 1997):

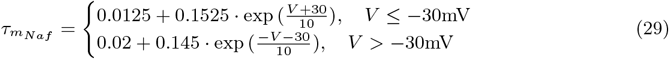

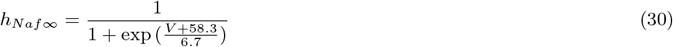

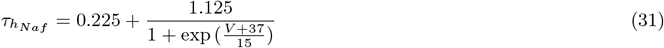

In TC and RE neurons, the kinetics for this current were based on dynamics described in Traub and Miles (Traub et al., 1991):

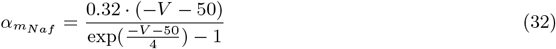

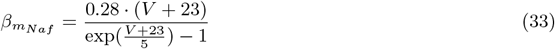

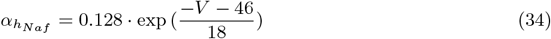

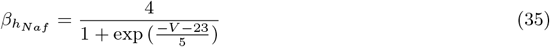

##### A.1.4 I_Nap_

The persistent, depolarization-activated *Na*^+^ current was described using kinetics presented by Traub et al. (Traub et al., 2003):

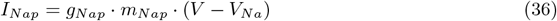

where *g*_*Nap*_ is the maximal conductance, and *m*_*Nap*_ is the activation variable. For all cortical neurons, the steady state activation function and decay time constant were described by the following:

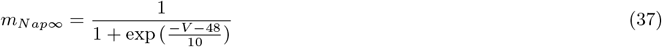

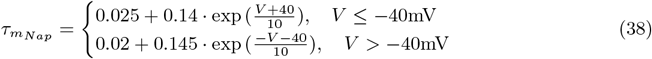

##### A.1.5 I_K_

The fast *K*^+^ current in both thalamic cells was described using the following equation:

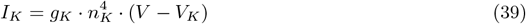

where *g*_*K*_ is the maximal conductance, *n*_*K*_ is the inactivation variable. All functions were based on dynamics described in Traub et al. (Traub et al., 1991):

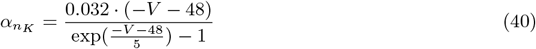

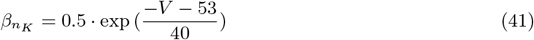

##### A.1.6 I_Kdr_

The delayed rectifier *K*^+^ current was described using kinetics presented by Traub et al. (Traub et al., 2003):

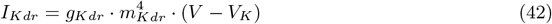

where *g*_*Kdr*_ is the maximal conductance and *m*_*Kdr*_ is the activation variable.

In neurons of the RS, IB and NRS type, the steady state activation function was described by the following:

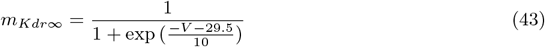

In LTS neurons, the steady activation function was defined as:

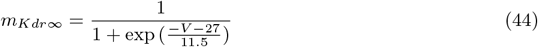

The time constant for all cortical neurons was described by the following:

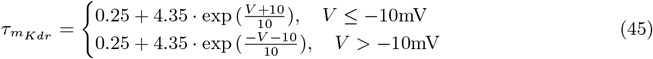

##### A.1.7 I_Ka_

The transient inactivating *K*^+^ current was modelled in all cortical cells using the following equation:

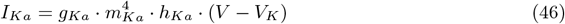

where *g*_*Ka*_ is the maximal conductance, and *m*_*Ka*_ and *h*_*Ka*_ are the activation and inactivation variables, respectively. The kinetics were based on dynamics described by Huguenard and McCormick (Huguenard and McCormick, 1992):

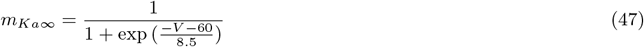

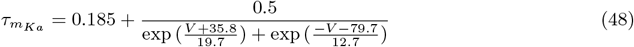

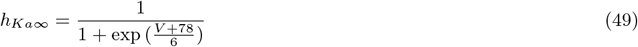

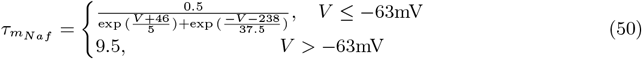

##### A.1.8 I_K2_

The slowly activating and inactivating *K*^+^ current equations followed Huguenard and McCormick (Huguenard and McCormick, 1992) and McCormick and Huguenard (McCormick and Huguenard, 1992):

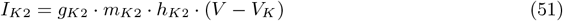

where *g*_*K*2_ is the maximal conductance, and *m*_*K*2_ and *h*_*K*2_ are the activation and inactivation variables, respectively. The kinetics for all cortical neurons were described by the following equations:

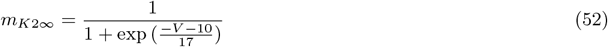

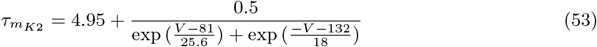

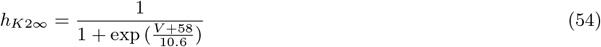

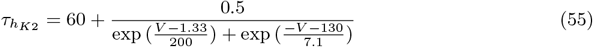

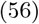

##### A.1.9 I_Km_

The non-inactivating, slow voltage-dependent *K*^+^ current was modelled using the following equation:

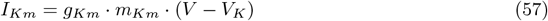

where *g*_*Km*_ is the maximal conductance, and *m*_*Km*_ is the activation variable.

In all cortical neurons, kinetics were based on dynamics presented by Traub et al. (Traub et al., 2003):

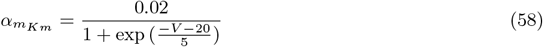

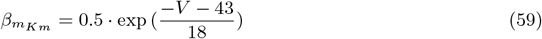

In TC and RE neurons, the kinetics were based on dynamics described in McCormick et al. (McCormick et al., 1993):

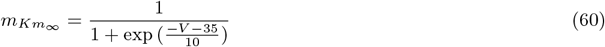

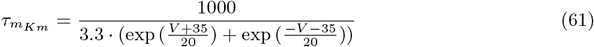

##### A.1.10 I_Kc_

The fast voltage and *Ca*^2+^-dependent *K*^+^ current was modelled using the equation presented by Traub et al. (Traub et al., 1994):

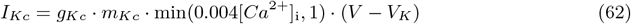

where *g*_*Kc*_ is the maximal conductance, *m*_*Kc*_ is the activation variable, and [*Ca*^2+^]_i_ dynamics are governed by dynamics described in A.2.

The forward and backward rate functions for all neurons of the RS, IB, and NRS type were based on kinetics described in Traub et al. (Traub et al., 2003):

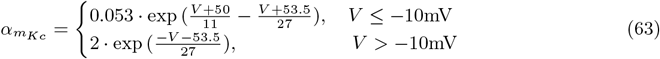

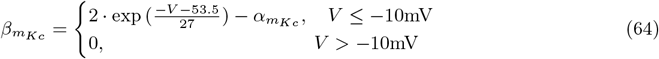

For LTS neurons, the forward and backward rate functions as described above were multiplied by a factor of 2.

##### A.1.11 I_Kahp_

The slow *Ca*^2+^-dependent *K*^+^ current responsible for afterhypolarization, was described using kinetics presented by Traub et al. (Traub et al., 1994):

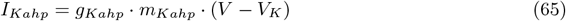

where *g*_*Kahp*_ is the maximal conductance, and *m*_*Kahp*_ is the activation variable. The forward rate function for all cortical neurons was dependent on the intracellular calcium dynamics described in Section A.2:

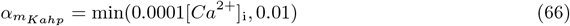

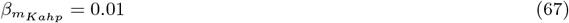

##### A.1.13 I_CaT_

The low-threshold *Ca*^2+^ current was modelled using the following equation:

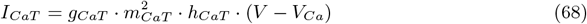

where *g*_*CaT*_ is the maximal conductance, *m*_*CaT*_ and *h*_*CaT*_ are the activation and inactivation gating variables, respectively. The dynamics were described differently for each thalamic neuron, as well as within the cortical cells.

For cortical neurons of the RS, IB, and NRS type, the kinetics were based on descriptions by Traub et al. (Traub et al., 2003):

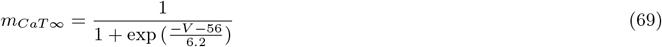

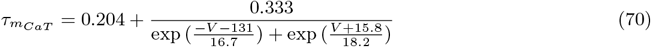

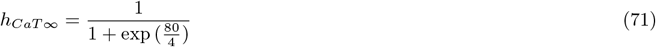

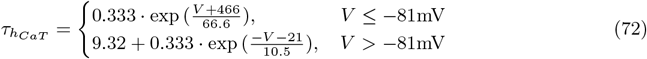

For the LTS interneuron, kinetics were based on descriptions by Destexhe et al. (Destexhe et al., 1996b):

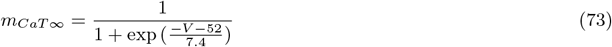

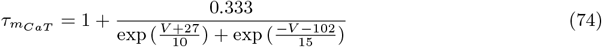

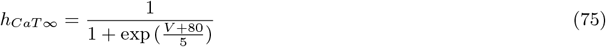

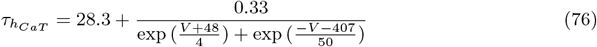

For RE neurons, kinetics were slightly modified from the description for LTS cells, as given by Traub et al. (Traub et al., 2005):

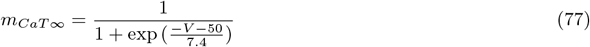

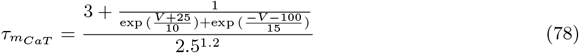

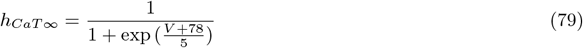

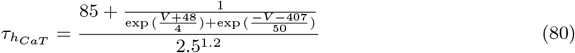

For TC neurons, activation *m*_*CaT*_ was taken to be at steady-state equal to *m*_*CaT \**_, and all steady-state activation and inactivation functions were based on the description by Huguenard and McCormick (Huguenard and McCormick, 1992), and the inactivation time constant was as described by Destexhe et al. (Destexhe et al., 1996a):

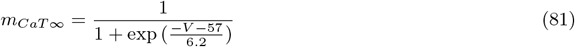

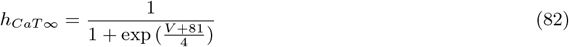

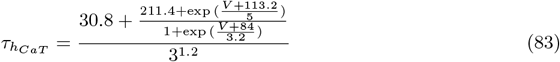

##### A.1.13 I_CaL_

The high-threshold *Ca*^2+^ current was modelled using the following equation:

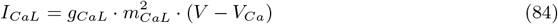

where *g*_*CaL*_ is the maximal conductance, and *m*_*CaL*_ is the activation variable. The forward and backward rate function for all cortical neurons were as described by Traub et al. (Traub et al., 2003):

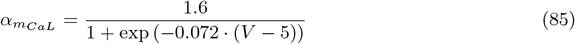

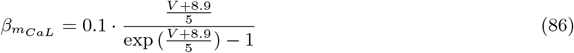

##### A.1.14 I_h_

The mixed *Na*^+^/*K*^+^ cation current was modelled differently for cortical cells and the TC cell. For all cortical cells, this was modelled using the following equation:

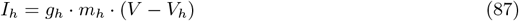

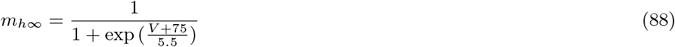

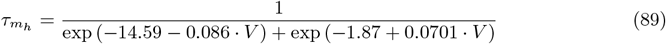

where *g*_*h*_ is the maximal conductance, and *m*_*h*_ is the voltage-dependent activation variable, based on the description by Huguenard and McCormick (Huguenard and McCormick, 1992):

For TC cells, in addition to being voltage-dependent, the current dynamics were also dependent on intracellular *Ca*^2+^ concentration:

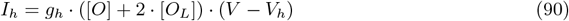

where *g*_*h*_ is the maximal conductance, and the activation variable, *m*_*h*_, is replaced by a weighted sum of channels in the open form and calcium-bound open form, based on the kinetic scheme described by Destexhe et al. (Destexhe et al., 1996a):

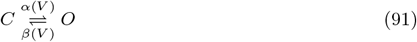

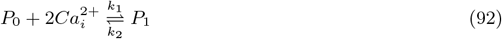

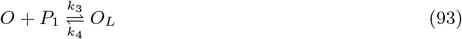

where voltage-dependent transitions between closed (*C*) and open (*O*) forms occur with rates described by *α*(*V*) and *β*(*V*), and intracellular *Ca*^2+^ ions bind to a regulating factor given by *P*_0_ (unbound) and *P*_1_ (bound) with rates of *k*_1_ = 2.5 *×*10^7^ mM^*\**4^ms^*\**1^ and *k*_2_ = 4 1*×*0^*\**4^ ms^*\**1^. The dynamics of *Ca*^2+^ ions is described in Section A.2. Furthermore, the open form of the channel binds with the calcium-bound regulator *P*_1_ to form *O*_*L*_ with rates *k*_3_ = 0.1 ms^*\**1^ and *k*_4_ = 0.001 ms^*\**1^.

The voltage-dependent transition rates between closed and open forms are described by:

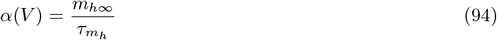

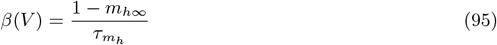

Where

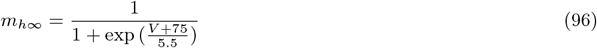

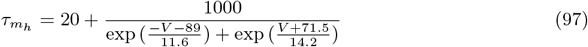

#### A.2 Calcium dynamics

Intracellular calcium dynamics were based on the model described by Destexhe et al. (Destexhe et al., 1993). A simple proportional model of *Ca*^2+^ diffusion was used with the change in calcium concentration described by:

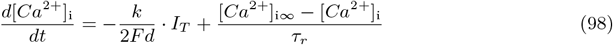

where [*Ca*^2+^]_i_ is measured in mM, *F* = 96489 Cmol^*\**1^ is Faraday’s constant, *d* denotes the depth at which calcium concentration was calculated beneath the membrane, *k*= 10^*\**4^ is a dimensionless unit conversion factor, *I*_*T*_ is the calcium current measured in mA/cm^2^, [*Ca*^2+^]_i*\**_ = 2.4 *×*10^*\**4^ mM is the steady state calcium concentration, and *τ*_*r*_ is the decay time constant. The depth at which the calcium concentration was calculated was 12 *×*10^*\**3^*µ*m for the IB and RS neuron, 4 *×*10^*\**3^*µ*m for the NRS neuron, 2 *×*10^*\**4^*µ*m for the LTS neuron, and 1*µ*m for both the TC and RE neurons. The decay time constants were 100ms for cortical cells of the IB, RS, and NRS type, 50 ms for the LTS type, and 5 ms for both TC and RE cells.

##### A.2.1 Equilibrium potential for *Ca*^2+^

The equilibrium potential for calcium, *V*_*Ca*_ was calculated according to the Nernst relation described in (Destexhe et al., 1993):

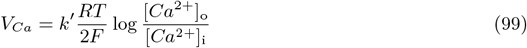

where *R* = 8.31 J mol^*\**1^K^*\**1^, *T* = 309.15^*°*^K, *F* = 96489 Cmol^*\**1^ is Faraday’s constant, [*Ca*^2+^]_°_and [*Ca*^2+^]_i_ denote extracellular and intracellular calcium concentrations respectively, and *k*^*′*^ = 1000 is a unit conversation factor for *V*_*Ca*_ in mV.

#### A.3 Synaptic dynamics

Synaptic currents were modelled in a manner similar to ionic currents as given by the following equation:

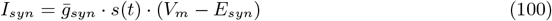

where 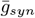 is the maximal synaptic conductance, *s*(*t*) is the synaptic conductance function and *E*_*syn*_ is the reversal potential of the synapse. The time-dependent conductance *s*(*t*) can be modelled in several ways depending on the type of neurotransmitter-mediated synapse, as described below (Destexhe et al., 1998b).

##### A.3.1 AMPA/GABAa-mediated synapse modelling

In the formalism by Destexhe et al. (Destexhe et al., 1994, 1998b), postsynaptic currents mediated by glutamate AMPA and GABAergic GABAa receptors are modelled using a gating variable *s*(*t*) which denotes the fraction of open synaptic channels at time *t*. Open synaptic channels refer to postsynaptic receptors that neurotransmitter molecules bind to, followed by the opening of ion channels. In this way, the synaptic conductance function is described by:

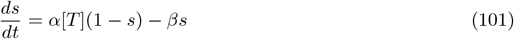

where [*T* ] represents the neurotransmitter concentration released into the synaptic cleft upon the arrival of a presynaptic spike, and *α* and *β* are rate constants describing the binding of neurotransmitter.

Upon the arrival of a presynaptic spike at *t* = *t*_0_, [*T* ] jumps to a maximal concentration value *C*_*max*_, and at *t* = *C*_*dur*_ (the duration of neurotransmitter-mediated pulse), [*T* ] falls back to 0. After the neurotransmitter-mediated pulse is gone, *s*(*t*) decays exponentially. Thus, solving Equation (101), we get:

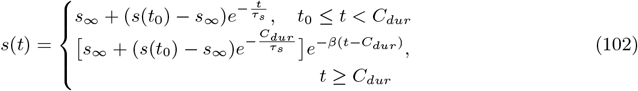

Where

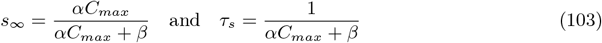

For AMPA-mediated synapses, *E*_*syn*_ = 0 mV, *C*_*max*_ = 0.5 mM and *C*_*dur*_ = 0.3 ms, while *α* and *β* parameters vary according to the synapse type, as presented in Tables 3–4. For GABAamediated synapses, *E*_*syn*_ =*\** 85 mV, *C*_*max*_ = 0.5 mM, and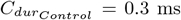, while values of *α*_*Control*_ and *β*_*Control*_ vary according to the synapse type, as presented in Tables 3–4. To model postALLO GABAa-mediated synapses, *α*_*post ALLO*_ = *α*_*Control*_ 1.58, *β*_*post ALLO*_ = *β*_*Control*_ 0.74, and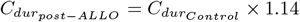.

**Table 3:** Forward (binding) rate, *α* parameters (in (ms mM)^*\**1^), for each synapse type, chosen to produce a default network state exhibiting spindle oscillations. Note that *α* values for inhibitory synapses shown here correspond only to GABAa synapses and reflect binding rates under Control receptor activity.

| Pre-synaptic neuron type | Post-synaptic neuron type |  |  |  |  |  |
| --- | --- | --- | --- | --- | --- | --- |
|  | RS | IB | NRS | LTS | TC | RE |
| RS | 0.05 | 0.05 | 0.1 | 0.5 | - | - |
| IB | 0.1 | 0.1 | 0.1 | 0.05 | - | - |
| NRS | 0.05 | 0.1 | 0.05 | 0.625 | 0.75 | 0.8 |
| LTS | 0.1 | 0.05 | 0.1 | - | - | - |
| TC | 1 | 1 | 0.07 | 0.149 | - | 0.94 |
| RE | - | - | - | - | 20 | 20 |

**Table 4:** Backward (unbinding) rate, *β* parameters (in ms^*\**1^) for each synapse type, chosen to produce a default network state exhibiting spindle oscillations. Note that *β* values for inhibitory synapses presented here correspond only to GABAa synapses and reflect unbinding rates under Control receptor activity.

| Pre-synaptic<br>neuron type | Post-synaptic neuron type |  |  |  |  |  |
| --- | --- | --- | --- | --- | --- | --- |
|  | RS | IB | NRS | LTS | TC | RE |
| RS | 0.15 | 0.05 | 0.01 | 0.2 | - | - |
| IB | 0.15 | 0.05 | 0.01 | 0.08 | - | - |
| NRS | 0.05 | 0.01 | 0.03 | 0.109 | 0.3 | 0.09 |
| LTS | 0.2 | 0.7 | 0.01 | - | - | - |
| TC | 0.15 | 0.15 | 0.006 | 0.048 | - | 0.18 |
| RE | - | - | - | - | 0.162 | 0.162 |

##### A.3.2 GABAb-mediated synapse modelling

The dynamics of GABAb-mediated synapses followed the formalism by Destexhe et al. (Destexhe et al., 1998b). GABAb-mediated synapses active *indirectly*, following a series of intracellular events. In particular, the neurotransmitter binding event activates an intracellular complex called a Gprotein which activates potassium channels ultimately hyperpolarizing and causing inhibition of the postsynaptic neuron. In this way, the synaptic conductance function is described by:

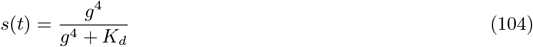

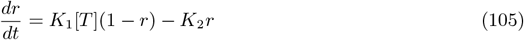

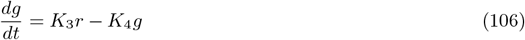

where *r* is the fraction of GABAb receptors in the activated form, *g* is the normalized G-protein concentration in activated form given in *µ*M, *K*_*d*_ = 100 *µ*M^4^ is the dissociation constant of G-protein binding on potassium channels, *K*_1_ = 0.09 (ms mM)^*\**1^ is the forward binding rate to the receptor, *K*_2_ = 0.0012 ms^*\**1^ is the unbinding rate, *K*_3_ = 0.18 ms^*\**1^ is the rate of G-protein production, *K*_4_ = 0.034 ms^*\**1^ is the rate of G-protein decay, and *E*_*syn*_ = *\**95 mV.

## Appendix

### B Model parameters and extended methodological details

#### B.1 Allopregnanolone application experiment curve fitting

Inhibitory postsynaptic currents mediated by GABAa receptors post application of allopregnanolone, obtained from experimental works (thereby referred to as *I*_*data*_), were fit using the following equation:

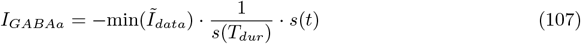

where

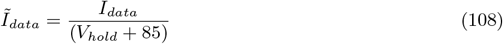

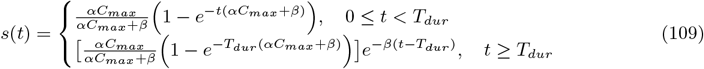

**Table B2:** Maximal conductance parameters (in mS/*cm*^2^) for each neuron type.

| $g_{ion}$ | RS | IB | NRS | LTS | TC | RE |
| --- | --- | --- | --- | --- | --- | --- |
| $g_{Na_f}$ | 200 | 200 | 200 | 170 | 90 | 200 |
| $g_{Na_p}$ | 0.16 | 0.16 | 0.1 | 0.16 | - | - |
| $g_K$ | - | - | - | - | 10 | 20 |
| $g_{K_{dr}}$ | 170 | 170 | 170 | 100 | - | - |
| $g_{K_a}$ | 75 | 17 | 20 | 1 | - | - |
| $g_{K_2}$ | 35 | 27 | 40 | 9 | - | - |
| $g_{K_m}$ | 43 | 40 | 55 | 35 | - | - |
| $g_{K_c}$ | 20 | 20 | 15 | 15 | - | - |
| $g_{K_{ahp}}$ | 0.5 | 3.5 | 35 | 0.1 | - | - |
| $g_{CaT}$ | 0.1 | 0.1 | 0.1 | 0.05 | 2 | 3 |
| $g_{CaL}$ | 10 | 11 | 2 | 0.1 | - | - |
| $g_h$ | 0.1 | 0.1 | 0.1 | 0.1 | $\mathcal{N}(0.0175, (8 \times 10^{-7})^2)$ | - |
| $g_{Leak}$ | 2 | 2 | 2 | 2.4 | 0.01 | 0.05 |
| $g_{K_{Leak}}$ | - | - | - | - | $\mathcal{N}(4, (3 \times 10^{-4})^2)$ | - |

**Table B3:**
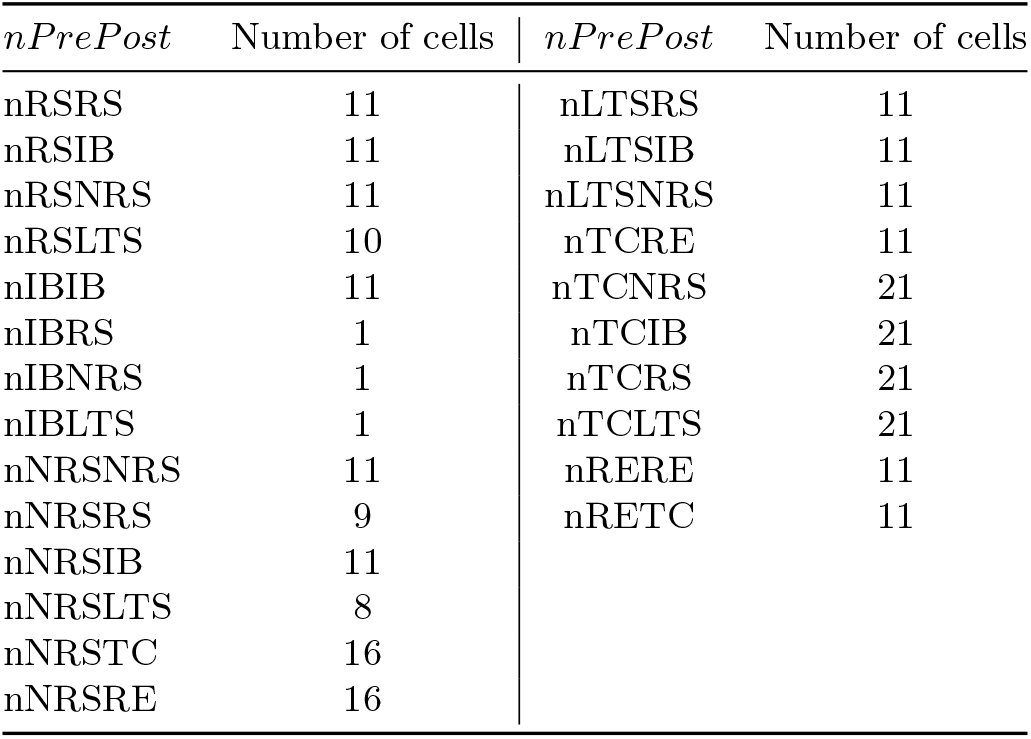
The number of presynaptic neurons that each postsynaptic neuron connects to (*nPrePost*) in the 5-95 (nIB:nRS) model configuration.

**Table B4:** The number of presynaptic neurons that each postsynaptic neuron connects to (*nPrePost*) in the 95-5 (nIB:nRS) model configuration.

| $nPrePost$ | Number of cells | $nPrePost$ | Number of cells |
| --- | --- | --- | --- |
| nRSRS | 11 | nLTSRS | 11 |
| nRSIB | 1 | nLTSIB | 11 |
| nRSNRS | 1 | nLTSNRS | 11 |
| nRSLTS | 1 | nTCRE | 11 |
| nIBIB | 11 | nTCNRS | 21 |
| nIBRS | 11 | nTCIB | 21 |
| nIBNRS | 11 | nTCRS | 21 |
| nIBLTS | 10 | nTCLTS | 21 |
| nNRSNRS | 11 | nRERE | 11 |
| nNRSRS | 11 | nRETC | 11 |
| nNRSIB | 9 |  |  |
| nNRSLTS | 8 |  |  |
| nNRSTC | 16 |  |  |
| nNRSRE | 16 |  |  |

All experimental data were scaled by factoring out the voltage dependent term for curve fitting purposes. This allowed for the isolation of the synaptic gating variable *s*(*t*) dynamics, making the analysis of the synaptic gating more direct and interpretable. We interpret the scaling factor min 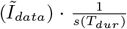 as the channel’s maximal intrinsic conductance where *s*(*T*_*dur*_) is equal to the maximum of the gating variable and min(*Ĩ*_*data*_) represents the maximal conductance.

We used MATLAB’s *lsqcurvefit* function to find the best-fitting *α, β* and *T*_*dur*_ parameters in Equation (107), for the given data. By pre-scaling the data, these parameters could be estimated to directly fit *s*(*t*). As illustrated in Figure B1, without pre-scaling, the relative importance of fitting different parts of the curve changes. This results in the optimization converging to different (non-ideal) local minima in the parameter space.

**Figure B1:**
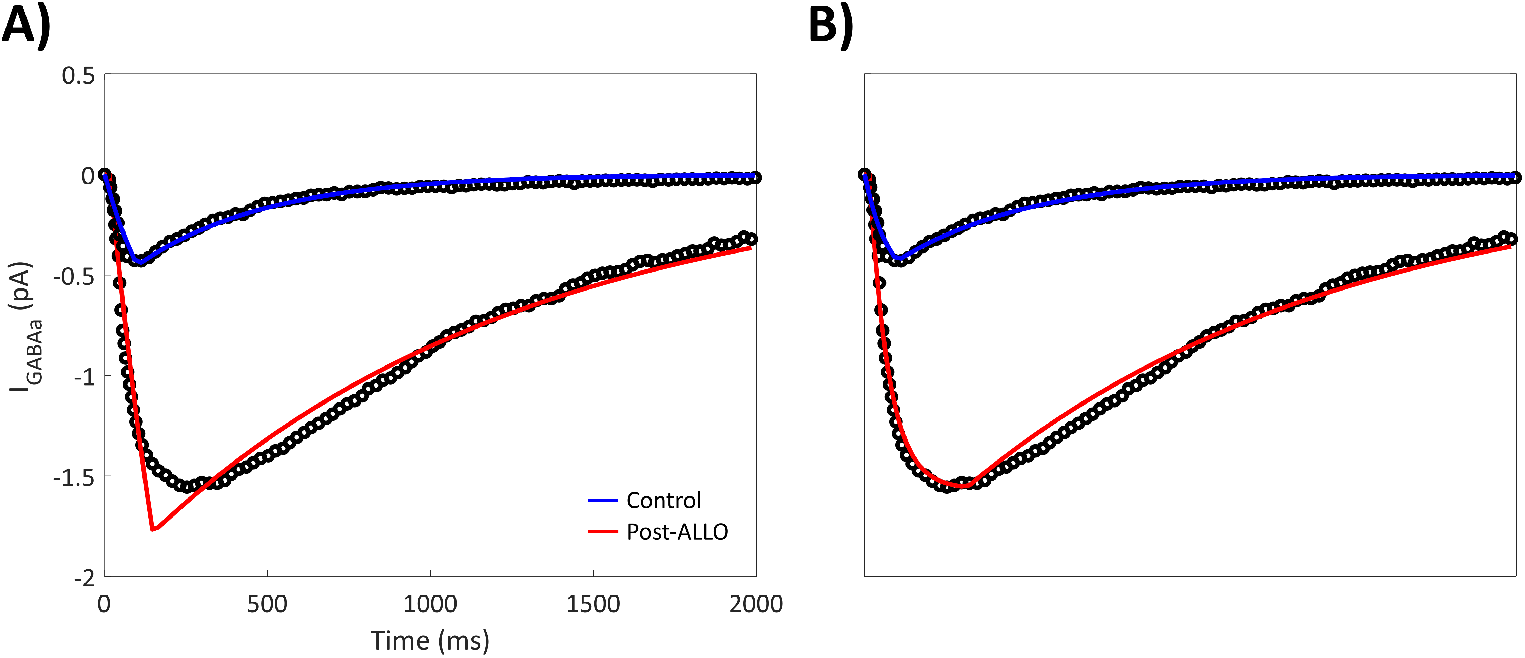
Sample curve fits of inhibitory postsynaptic currents post application of allopregnanolone, without (A) and with (B) pre-scaling data by factoring out the voltage dependent term.

**Figure B2:**
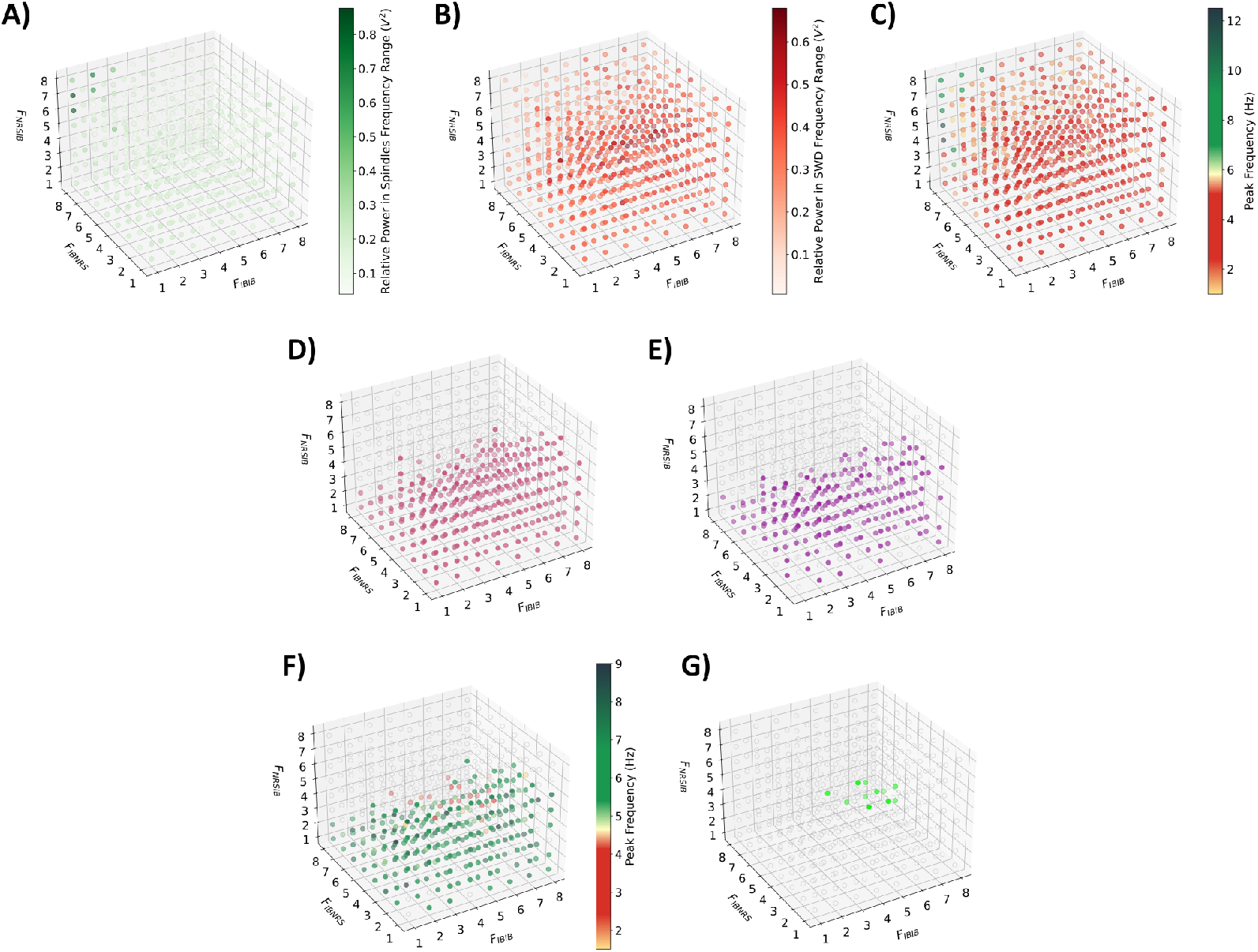
3D visualization of the grid search results across 8^3^ = 512 parameter combinations within the (*F*_*IBIB*_, *F*_*IBNRS*_, *F*_*NRSIB*_) parameter space using the 50-50 (nIB:nRS) model configuration. (A–C): Initial search using Control synapses with cortical GABAa conductance at 10% baseline, were filtered by examining the magnitude of: (A): relative power within spindles range, (B): relative power within SWDs range, and (C): peak frequency from power spectral density analysis. (D): Filtered parameter combinations from the initial search. (E): Parameter combinations that produce spindle-like oscillations when simulated with cortical GABAa conductance restored to normal (100% baseline) levels. (F): Parameter combinations that produce continued SWDs (in red) when the network is simulated using 10% baseline cortical GABAa conductance and post-ALLO synapses. (G): Final parameter set of (*F*_*IBIB*_, *F*_*IBNRS*_, *F*_*NRSIB*_) for which the 50-50 (nIB:nRS) model exhibits continued SWDs post application of ALLO.

#### B.2 Modelling increased frontocortical connectivity

